# CDKL5 phosphorylates neuronal ELAVL proteins to promote mRNA binding, protein synthesis and visual cortex development

**DOI:** 10.64898/2026.04.07.716766

**Authors:** Simeon R. Mihaylov, André T. Lopes, Margaux Silvestre, Gaia Bianchini, Helen R. Flynn, Almaz Huseynova, Stephanie Strohbuecker, Llywelyn Griffith, Cristina Militti, Lucas L. Baltussen, Xiuming Yuan, Gabriel Morel, Suzanne Claxton, Kelvin Dempster, Flora C. Y. Lee, Oguz Kanca, Marcel Köhn, Mark Skehel, Jernej Ule, Florencia Iacaruso, Sila K. Ultanir

## Abstract

Loss-of-function mutations in the X-linked CDKL5 gene lead to a severe neurodevelopmental disorder characterized by early-onset epilepsy, known as CDKL5 Deficiency Disorder (CDD). Despite its clinical significance, the physiological substrates of the serine/threonine kinase CDKL5 and its roles in neuronal development remain poorly understood. To address this, we performed quantitative phosphoproteomics analysis in *Cdkl5* knockout (KO) mouse brains, identifying 22 CDKL5 substrates involved in diverse cellular functions. Among these, we focused on the neuronal RNA-binding proteins (nELAVLs) ELAVL2, ELAVL3, and ELAVL4, as these represented the only evolutionarily conserved phosphorylation and are known regulators of neuronal differentiation. Through kinase assays and individual-nucleotide resolution crosslinking and immunoprecipitation (iCLIP), we found that CDKL5 phosphorylates S119/131 in ELAVL2/3/4, promoting their cytoplasmic localization and enhancing their binding to target mRNAs at 3’UTRs. Loss of CDKL5 activity in neurons caused reduced new protein synthesis, as measured by puromycin incorporation; this phenotype was rescued by knockdown of the nELAVL inhibitor long non-coding RNA, RNY3, revealing an essential function of CDKL5 in enhancing protein synthesis via nELAVL phosphorylation. To investigate the *in vivo* functions of nELAVL phosphorylations, we generated *Elavl2/3/4* phosphomutant mice and found that collectively nELAVL phosphorylations are required for viability. Proteomic and transcriptomic analyses of *Elavl2/3* homozygous phosphomutants, which exhibited sub-viability, revealed compensatory upregulation of ELAVL4 and synaptic proteins. Functionally, in *Elavl2/3/4* triple heterozygous mice Neuropixels recordings in the primary visual cortex showed deficits in receptive field properties and orientation tuning, revealing the role of nELAVL phosphorylation for accurate cortical circuit formation. Our study uncovers a crucial role for CDKL5 in regulating nELAVL-mediated protein synthesis and the development of cortical circuits.

## Introduction

Cyclin-dependent kinase-like 5 (CDKL5) is a serine/threonine kinase in the CMGC family of kinases ^1^. Loss-of-function mutations in the X-linked *CDKL5* lead to a rare neurodevelopmental disorder known as CDKL5 deficiency disorder (CDD, OMIM 300672, 300203) ^2, 3, 4, 5^. CDD is characterized by early-onset intractable epilepsy, hypotonia, vision and speech impairments, as well as profound neurodevelopmental impairments ^6, 7^. Missense mutations are found almost exclusively in the kinase domain of CDKL5, some of which reduce kinase activity^8^ implicating a critical role for its kinase activity in CDD pathology^7, 9, 10, 11^. The prevalence of CDD is estimated at 1 in 42,000 births, making it one of the most common forms of genetic epilepsy ^6, 12^. Patients with CDKL5 deficiency disorder need lifelong care, and there are currently no targeted therapies available. Existing treatments primarily consist of anti-epileptic medications.

Widespread CDKL5 expression in brain starts at late embryonic stages, peaks during postnatal stages and remains high in adults in rodents and follows a similar trajectory in humans ^13, 14, 15, 16^. The cellular and molecular roles of CDKL5 have been investigated in animal models, demonstrating that CDKL5 plays roles in regulating the microtubule cytoskeleton, neuronal differentiation, excitability, and synaptic function^3, 17, 18, 19, 20, 21, 22, 23, 24, 25, 26^. To fully understand its function, it is critical to uncover substrates of CDKL5. Although several CDKL5 substrates have been documented in brain^8, 13, 27^ and other cell types^28, 29, 30^, the full range of CDKL5 substrates has not yet been established, and the critical substrates required for brain development remains unidentified.

ELAVL proteins are a family of highly conserved RNA binding proteins implicated in a variety of RNA processes, including splicing, polyadenylation, transport, stability and protein translation ^31, 32, 33, 34, 35, 36, 37, 38, 39^. The family consists of four members: ELAVL1, ELAVL2, ELAVL3 and ELAVL4 (HuR, HuB, HuC and HuD, respectively) all of which contain three highly conserved RNA recognition motifs (RRMs), two at the N-terminus and one at the C-terminus. RRM1 and RRM2 are crucial for the binding of mRNA while RRM3 is important for dimerization and interactions with other proteins^40, 41, 42, 43, 44, 45^. While ELAVL1 is ubiquitously expressed across tissues ^46^, ELAVL2/3/4, also known as neuronal ELAVLs (nELAVLs) are mostly expressed in the brain ^47, 48, 49, 50^. nELAVLs bind to a diverse array of mRNA targets that were identified in both mouse and human ^51, 52^. These targets include genes required for neuronal differentiation, highlighting nELAVL proteins’ significant role in neurodevelopment. However, the regulation of ELAVL proteins by phosphorylation during development is not well understood.

In this paper we used global phosphoproteomics approach to determine the substrates of CDKL5 in the mouse brain. We confirmed several known substrates and identified 16 novel substrates. Among these, we validated the CDKL5-dependent phosphorylation of nELAVL proteins at the highly conserved Ser119 in ELAVL2, ELAVL3 and Ser131 in ELAVL4, located between the first and the second RNA binding domains. Our findings in *Cdkl5* KO and *Elavl2/3/4* phosphomutant mouse models indicate that nELAVL phosphorylation is necessary for nELAVLs’ cytoplasmic localization, binding to numerous mRNA 3’UTRs and enabling efficient mRNA translation in neurons. Our findings demonstrate that ELAV plays a vital role during development, and its disruption leads to impairments in visual processing in adult mice.

## Results

### Phosphoproteomic analysis of *Cdkl5* knockout mice uncovers novel substrates

To uncover the changes in phosphorylation in the brains of *Cdkl5* knockout (KO) mice ^53^, we used global quantitative proteomics and phosphoproteomics analyses with TMT isobaric labeling **(Fig. 1a)**. We compared whole-hemisphere protein extracts from 5 wild-type (WT) and 5 *Cdkl5* KO male littermates at postnatal day 10 (P10). Among 7,625 proteins we identified and quantified in this experiment, we observed 8 proteins to be increased and 14, including CDKL5, to be reduced in C*dkl5* KO brains *(limma* p-value<0.05 and log2 fold change (lfc)>0.2) **(Fig. 1b and Supplementary Data 1)**. We next analyzed phosphoprotein levels to identify novel CDKL5 substrates and we detected close to 30,000 phosphosites finding several significantly changed sites in *Cdkl5* KOs **(Fig. 1c and Supplementary Data 1)**. Phosphopeptides which belong to CDKL5 were reduced due to the absence of CDKL5 protein, as expected **(Fig. 1c, blue)**. We found that there were 24 phosphorylation sites, corresponding to 22 proteins, which were significantly reduced in *Cdkl5* KOs (*limma* p-value<0.05, lfc>0.2), strikingly containing the CDKL5 consensus motif RPXS* ^13, 54^ **(Fig. 1c, pink; Table 1 and Supplementary Data 1)**. Among these were known CDKL5 substrates MAPRE2/EB2, ARHGEF2, MAP1S ^8, 13^, Dlg5 ^8^, EP400 ^28^ and CACNA1E/Cav2.3 ^27^, validating our assay. In addition, 16 novel candidate substrates were identified implicating CDKL5 in a range of molecular functions including synaptic transmission (e.g. Syngap1, Grin2b, Shank1), gene regulation (e.g. Bcl11a, Bcl11b, GlyR1) and mRNA binding (e.g. ELAVL2/3/4, Ralyl) **(Table 1)**. The observed minor changes in protein levels, which did not coincide with the identified candidate substrates, suggest that the reduction of substrate phosphorylation is not due to a reduction of total protein levels. These findings highlight the extensive functions of CDKL5 in brain development.

**Fig. 1:**
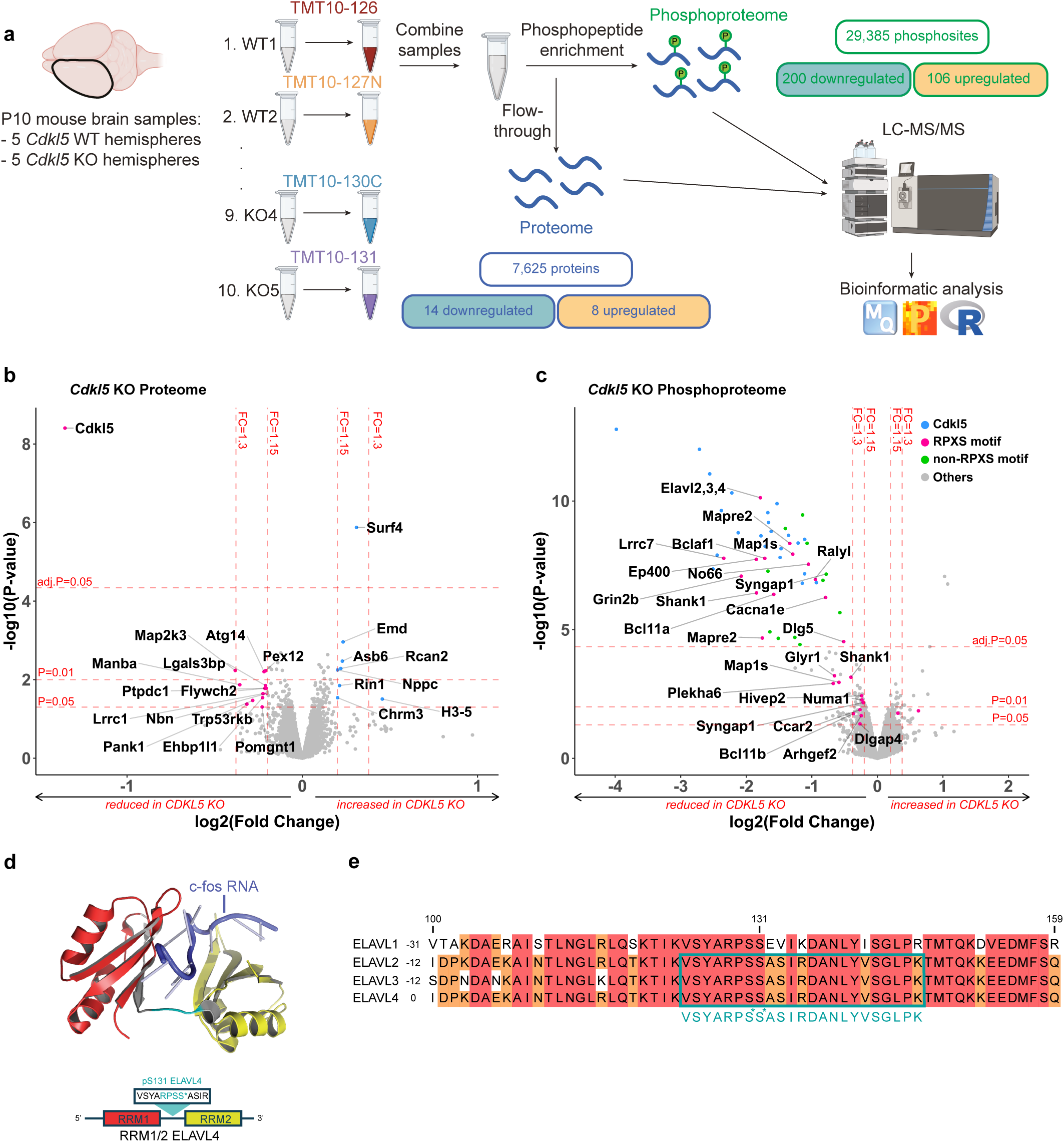
Novel substrates of CDKL5 in mouse brain. **a** Schematic of (phospho)proteomics approach using Tandem Mass Tag (TMT) mass spectrometry to label samples for accurate multiplexing quantification of phosphoprotein and total protein levels. In this experiment, hemispheres from 5 *Cdkl5* WT and 5 *Cdkl5* KO P10 mice were compared. **b** Volcano plot of total protein levels comparison between *Cdkl5* KO and WT mouse brains. Each point represents a different protein. Significance is defined as *limma* p<0.05 and log2 fold change (lfc)>0.2 = fold change (fc)>1.15. **c** Volcano plot of differential levels of phosphosites between *Cdkl5* KO and WT mouse brains. Each data point represents a different phosphosite. Phosphosites that belong to CDKL5 protein are shown in blue. Phosphosites that contain the RPXS* consensus motif and are significantly changed are shown in pink. Significantly reduced phosphosites that do not contain RPXS* are shown in green. Significance is defined as *limma* p<0.05 and log2 fold change (lfc)>0.2 = fold change (fc)>1.15. **d** Crystal structure (PDB:1FXL) representing RRM1 and RRM2 of human ELAVL4 binding to the AU-rich element of c-fos RNA. CDKL5 consensus sequence is highlighted in cyan with S* indicating the phosphorylated residue (pS131). **e** Protein sequence alignment of ELAVL1, 2, 3, and 4 (UniProt IDs: P70372, Q60899, Q60900 and Q61701, respectively) between amino acids 100 and 159 (mELAVL4) with green square indicating the ELAVL phosphopeptide that was detected and quantified by TMT-labelled mass spectrometry.

**Table 1.**
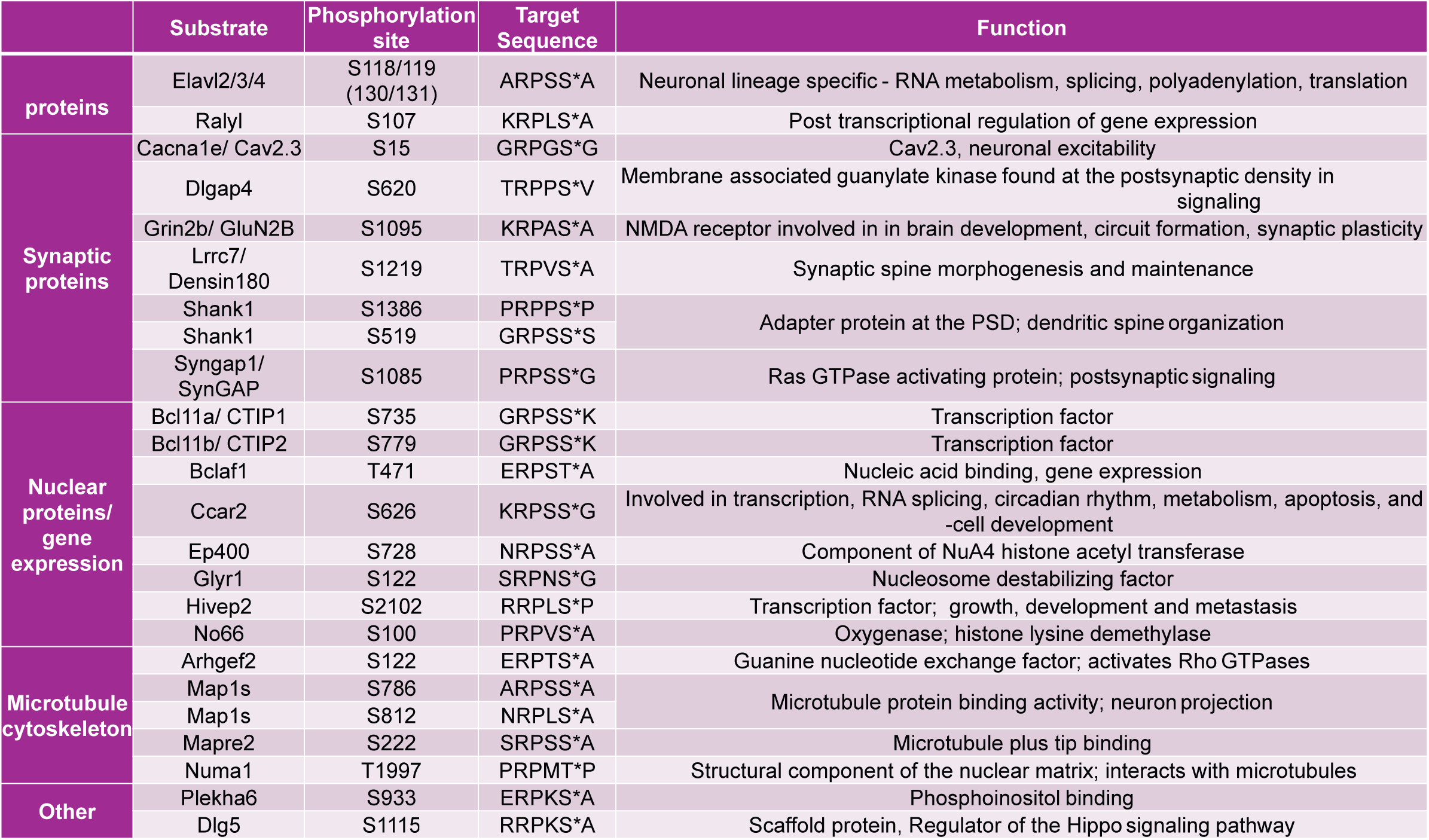
Phosphoproteomics screen identifies novel substrates of CDKL5. Phosphorylation sites containing the RPXS* CDKL5 consensus motif that are reduced in CDKL5 KO mice are included in this table. In addition to previously reported CDKL5 substrates (Cav2.3, Map1s, Mapre2/EB2, Arhgef2 and Dlg5), novel CDKL5 substrates are identified. Novel substrates include RNA binding proteins (e.g. Elavl2/3/4, Ralyl, synaptic proteins (e.g. Shank1, Grin2b), transcription factors (Bcl11a, Bcl11b) and microtubule binding protein (e.g. Numa1). The CDKL5 target phosphorylation site in mice and known functions of these proteins are shown.

### CDKL5 phosphorylates neuronal ELAVL (nELAVL) proteins between RRM1 and RRM2 in mice and humans

Neuronal ELAVL (ELAVL2/3/4) proteins have conserved functions in neuronal differentiation ^37, 52^. Interestingly, out of all the substrates we identified **(Table 1)**, nELAVLs were the only ones with phosphorylation sites conserved in Drosophila orthologs, pointing to an important functional role **(Supplementary Data 2)**. We thus focused our attention to the role of CDKL5’s phosphorylation of nELAVLs. The identified CDKL5-dependent phosphorylation at Serine 119 (pS119) ELAVL2/3 and Serine 131 (pS131) ELAVL4 is located between the RNA recognition motif 1 (RRM1) and RRM2 **(Fig. 1d)**. Since ELAVL2/3/4 are highly conserved, the phosphorylated peptide detected in our mass spectrometry spectra is identical in amino acids among the ELAVL2/3/4 family members **(Fig. 1d and Supplementary Fig. 1a)**. The localization probability of the pS in the peptide was 0.585 suggesting that either S118/130 or S119/131 could be phosphorylated **(Fig. 1e)**.

To confirm that nELAVLs are direct substrates of CDKL5, we performed *in vitro* kinase assays using purified recombinant CDKL5 kinase domain and a truncated ELAVL4 (RRM1-2) to represent nELAVL proteins, followed by Phos-tag^TM^ gel electrophoresis. The shift in molecular weight upon phosphorylation indicated that ELAVL4 is a direct substrate **(Fig. 2a)**. We next generated a phospho-specific antibody targeting the common RPXS* phosphopeptide pS119 ELAVL2/3 and pS131 ELAVL4 (anti-p-nELAVL). In HEK293T cells, we co-expressed HA-tagged human CDKL5 (hCDKL5) WT or kinase dead (KD) mutant with full-length GFP-tagged mouse ELAVL4 (mELAVL4) WT or phosphomutant Ser131Ala (SA) and probed with anti-p-nELAVL. We found that CDKL5 phosphorylates ELAVL4 at S131 and the antibody did not detect any signal in the absence of CDKL5 activity or in S131A mutant, confirming its specificity **(Fig. 2b)**. Given the high sequence identity within the linker region between RRM1 and RRM2 **(Fig. 1e)**, these findings show that CDKL5 can phosphorylate all nELAVLs *in vitro*. To validate nELAVLs as *in vivo* substrates, we performed Western blots in *Cdkl5* WT and KO mouse brain lysates and found a major reduction of endogenous p-nELAVL in KO brains **(Fig. 2c-f)**. ELAVL2, 3 and 4 have over 80% sequence homology and are 360, 367 and 385 amino acids long, respectively, with highly similar molecular weights. nELAVL antibody detected a band below 37 kDa, which corresponded to the molecular weight detected with p-nELAVL antibody **(Fig. 2c)**. We quantified the p-nELAVL/ total nELAVL ratio and found it to be diminished to approximately 15% of control levels in CDKL5 KOs **(Fig. 2d).** The remaining phosphorylation is likely due to other kinases such as CDKL2, known to phosphorylate ∼15% of the CDKL5 substrate EB2 ^55^. CDKL5 was absent and total nELAVL was unchanged in CDKL5 KOs **(Fig. 2e,f)**. Finally, we examined whether CDKL5-dependent regulation of phosphorylated nELAVL is conserved in human neurons. Using protein lysates from iPSC-derived neurons from 2 CDD patients and their control family members ^56^, we found that the normalized p-nELAVL levels to total nELAVL are significantly reduced in patients **(Fig. 2g-i)** whereas, despite a trend, total nELAVL levels were not significantly different showing a conserved regulation in humans. These results show that nELAVLs are substrates of CDKL5 in mouse brain and in human neurons.

**Fig. 2:**
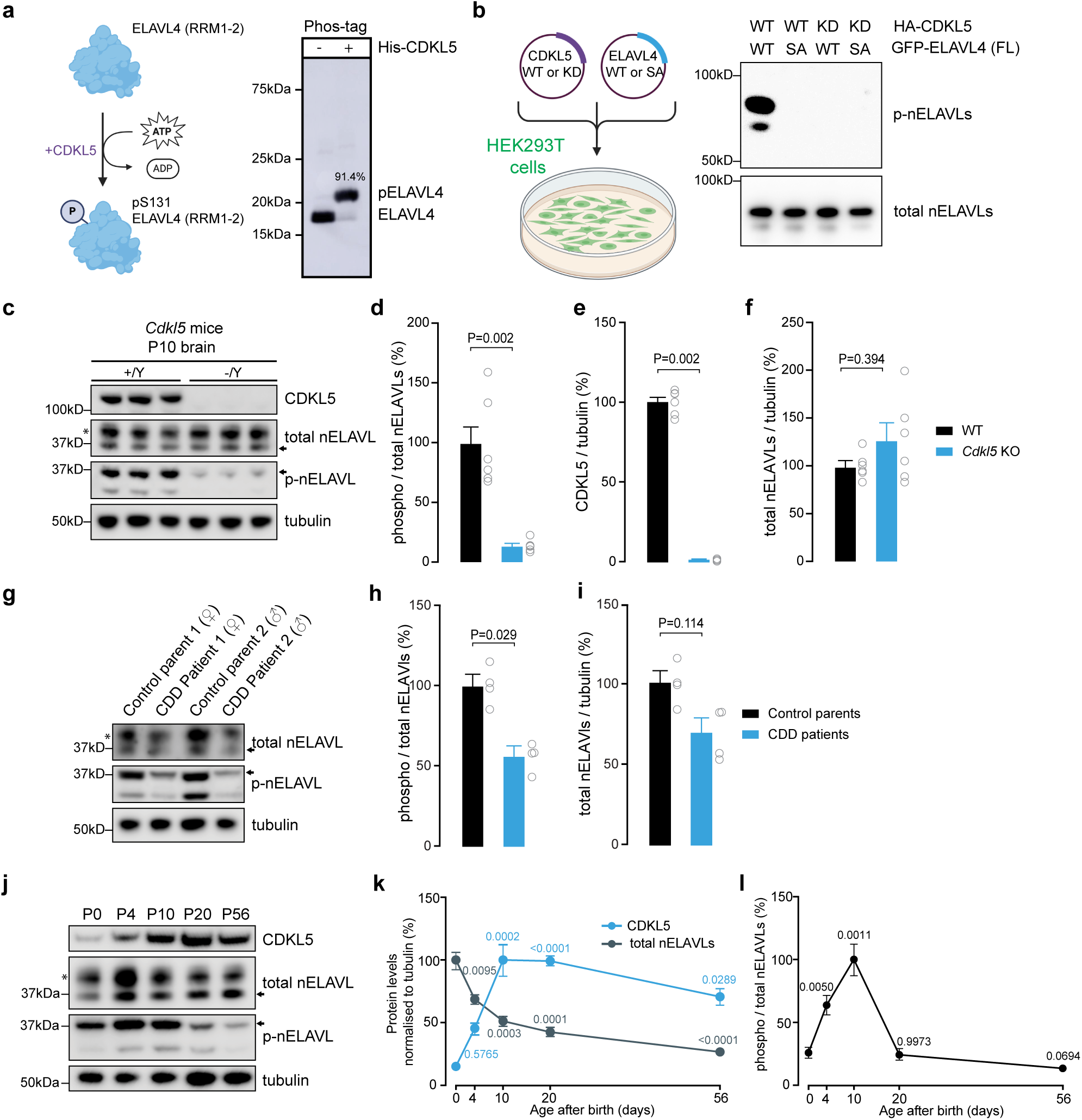
Neuronal ELAVL (nELAVLs) proteins are physiological substrates of CDKL5. **a** Schematic of an *in vitro* kinase assay reaction and Phos-tag^TM^ gel of the reaction showing unphosphorylated and phosphorylated ELAVL4. In the presence of His6-tagged CDKL5, ELAVL4 (RRM1-RRM2) was 91.4% phosphorylated. **b** Validation of phospho-specific antibody against Ser 131 in ELAVL4. HEK293T cells co-transfected with either wild-type (WT) or kinase dead mutant (KD) HA-tagged CDKL5 kinase domain and either WT or phosphomutant (SA; S131A) GFP-tagged full-length (FL) ELAVL4. Western blot shows specificity of the antibody against S131 when phosphorylated by CDKL5. **c** Western blots showing expression of CDKL5, total nELAVLs and tubulin and level of nELAVLs phosphorylation in P10 *Cdkl5* KO mouse brain. The arrows indicate the location of nELAVLs. **d-f** Quantification of nELAVLs phosphorylation, CDKL5 expression and total nELAVLs expression, respectively, in P10 *Cdkl5* full KO brain hemisphere. Two-sided Mann-Whitney test. n = 3 animals per genotype with 2 technical replicates. (In (**d**), WT=100, SEM=14.38, KO=13.83, SEM=1.827; in (**e**), WT=100, SEM=3.122, KO=0.8295, SEM=0.3386; in (**f**), WT=100, SEM=5.132, KO=44.01, SEM=17.97). **g** Western blots showing expression of total nELAVLs and level of nELAVLs phosphorylation and tubulin in iPSC-derived neurons from 2 CDD patients and their control parents. The arrows indicate the location of nELAVLs. **h, i** Quantification of nELAVLs phosphorylation and total nELAVLs expression, respectively. Two-sided Mann-Whitney test. n = 2 patients with 2 technical replicates (in (**h**), controls=100, SEM=7.605, patients=56.20, SEM=5.491; in (**i**), controls=100, SEM=8.126, patients=68.79, SEM=9.933). **j**, Western blots showing expression of CDKL5, total nELAVLs, tubulin and lever of nELAVLs phosphorylation in P0, P4, P10, P20 and P56 WT mouse brain. The arrows indicate the location of nELAVLs. **k** Quantification of total nELAVLs and CDKL5 expression. **l** Quantification of nELAVLs phosphorylation. Multiple comparisons test. n = 3 animals with 2 technical replicates. P-values compared with day 0 are shown on the graphs. Highest protein level is set at 100% (**k**), for CDKL5 Kruskal-Wallis test with Dunn’s post-hoc test P0=15.33, SEM=2.401, P4=45.52, SEM=4.751, P10=100, SEM=12.52, P20=99, SEM=3.444, P56=70.733, SEM=6.575; for total ELAVLs Brown-Forsythe and Welch ANOVA tests with Dunnett’s T3 post-hoc test P0=100, SEM=7.013, P4=68.66, SEM=3.505, P10=51.35, SEM=3.142, P20=42.57, SEM=2.649, P56=26.80, SEM=2.368; in (**l**), Brown-Forsythe and Welch ANOVA tests with Dunnett’s T3 post-hoc test P0=26.12, SEM=4.078, P4=63.82, SEM=7.822, P10=100, SEM=12.22, P20=24.45, SEM=4.302, P56=13.70, SEM=1.802). In (**c**), (**g**) and (**j**), * indicates a non specific band

CDKL5 expression increases during early postnatal development and peaks in adult, yet substrate phosphorylations are highest during early postnatal development ^13^, we next investigated if p-nELAVL levels are regulated developmentally after birth. In an age range from P0 to P56, we found that phosphorylation of nELAVL peaks at early postnatal development P4-P10 and subsequently decreases in adults **(Fig. 2j-l)**. This suggests that the CDKL5-mediated phosphorylation of nELAVLs plays a pivotal role in early postnatal stages, a crucial phase for brain development. Our results confirm that nELAVLs are *bona fide* substrates of CDKL5 and show that their phosphorylation is prominent during neuronal development.

### Phosphorylation promotes the cytoplasmic localization of neuronal ELAVLs

We next investigated the cellular role of nELAVL phosphorylation by CDKL5. The phosphorylation site for CDKL5 on nELAVL proteins is located between the first two RRMs which interact with RNA, as revealed by their crystal structure with c-fos **(Fig. 1d)** ^44^. We thus hypothesized that phosphorylation could influence nELAVLs’ RNA binding function. To test this, we conducted Bio-layer Interferometry (BLI) assay to compare the binding of phosphorylated and non-phosphorylated purified recombinant ELAVL4 (RRM1-2) to a U-rich RNA probe (biotin-U(x20)), which is known to be the primary binding motif for nELAVLs ^57^ **(Supplementary Fig. 2a-d)**. Additionally, a control GC-rich probe (biotin-GC(x10)) showed no binding, as expected **(Supplementary Fig. 2c)**. Our findings indicated that phosphorylated ELAVL4 binds to the U-rich probe with a 2:1 stoichiometry *in vitro*, with a primary equilibrium dissociation constant (KD) of 21 nM and a secondary KD of 6 μM **(Supplementary Fig. 2e)**. Non-phosphorylated ELAVL4 exhibited a similar binding model and dissociation constants (KD1 = 20 nM and KD2 = 4 μM) **(Supplementary Fig. 2f)**. These results show that the phosphorylation did not alter ELAVL4’s RNA binding *in vitro*, likely due to several other interactions between RRM1/2 and mRNA ^58^.

Phosphorylations often alter subcellular localization of proteins. To investigate if phosphorylation altered localizaion, we transfected NSC-34 cells with either wild-type (WT) or phosphomutant (S131A) full-length GFP-tagged ELAVL4. We found that S131A had increased nuclear/cytoplasmic ratio when compared to WT. Double phosphomutant S130/131A ELAVL4, showed an even greater increase in nuclear localization **(Fig. 3a,b)**, indicating that both of these two serine phosphorylations contribute to cytoplasmic localization. We next asked whether similar localization defects can be observed for endogenous nELAVLs in *Cdkl5* KO mouse brains. We stained P10 brain sections with an antibody targeting ELAVL3 and ELAVL4 and a second ELAVL2-specific antibody. We observed that ELAVL3/4 are present in most neurons while ELAVL2 had sparse expression in cortex **(Supplementary Fig. 3a-b)**, agreeing with previous findings ^50^. We observed an elevated nuclear-to-cytoplasmic ratio of ELAVL3/4 **(Fig. 3c,d)** and ELAVL2 **(Supplementary Fig. 3c,d)** in cortical layer 2/3 pyramidal neurons of *Cdkl5* KO brains. A comparable shift in ELAVL3/4 immunostaining was also detected in layer 5 neurons **(Supplementary Fig. 3e,f)** and in dentate gyrus granule cells **(Supplementary Fig. 3g,h)**, the latter of which exclusively express the neuronal isoform ELAVL3 ^52^. These findings demonstrate that nELAVL phosphorylation facilitates their cytoplasmic localization, and that endogenous ELAVL2 and ELAVL3 show defect in cytoplasmic localization in *Cdkl5* knockout models. These data suggest a disruption of nELAVL’s normal function in the absence of CDKL5-mediated phosphorylation.

**Fig. 3:**
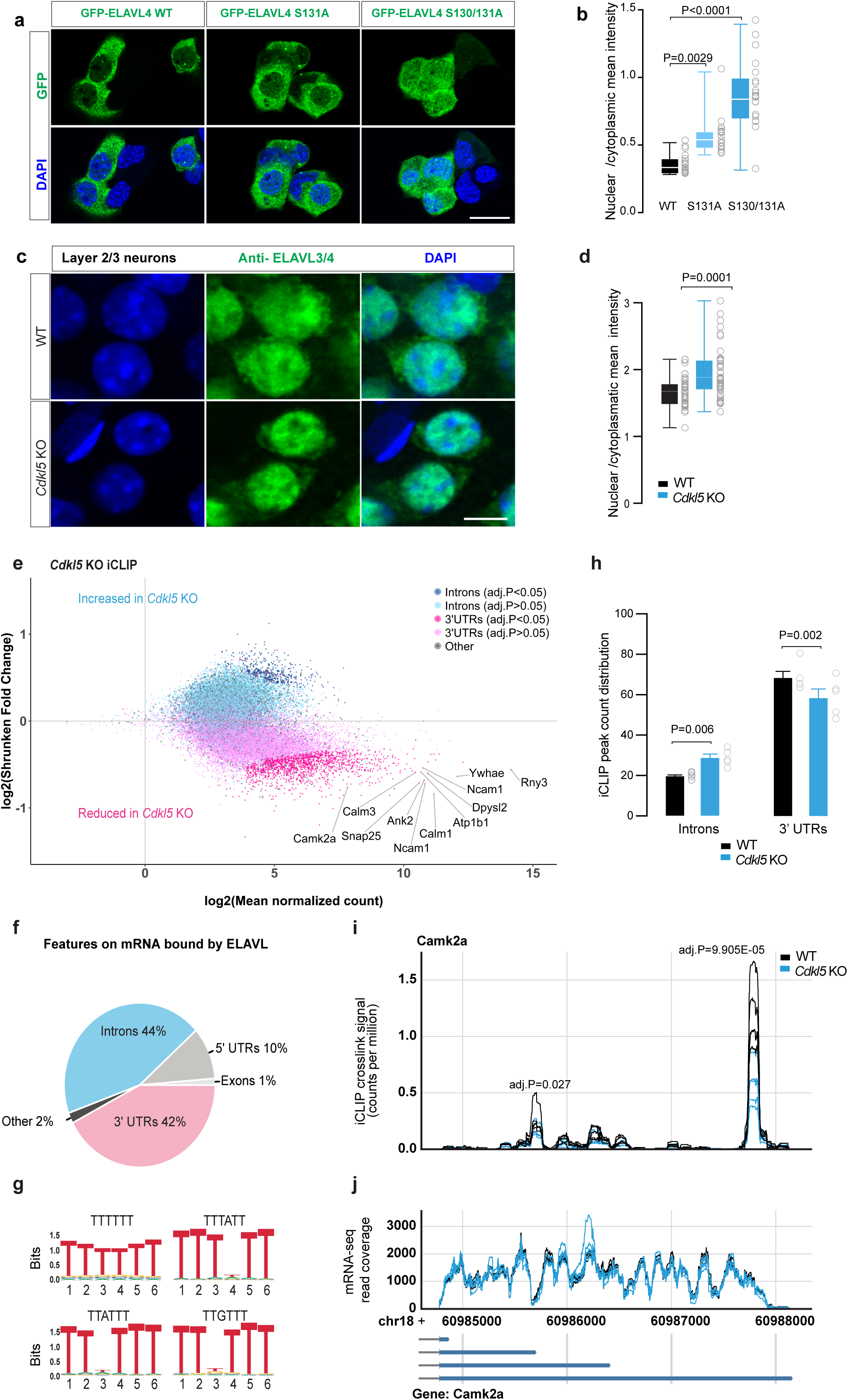
CDKL5 phosphorylation of nELAVLs enhances their cytoplasmic localization and mRNA 3’UTR binding. **a** NSC-34 cells transfected with either GFP-ELAVL4, GFP-ELAVL4 S131A or GFP-ELAVL4 S130/131A showing altered nuclear/cy-toplasmic localization. Scale bar = 20 μm. **b** Quantification of nuclear/cytosplasmatic mean GFP-ELAVL4 fluorescence ratio. n=16-20 cells/group. Ordinary one-way ANOVA was used to assess statistical significance (WT=0.3526, SEM=0.01730; S131A=0.5620, SEM=0.03601; S130/131A=0.8685, SEM=0.05536). **c** Layer 2/3 pyramidal neurons from WT and *Cdkl5* knockout mice at P10 stained with an antibody raised against ELAVL3/4. The scale bar is 10 μm. **d** Quantification of ELAVL3/4 immunofluo-rescence in nuclear/cytosplasmatic location in cell body. n=39-41 neurons/group. (Two-sided unpaired t-test, WT=1.460, SEM=0.03033; *Cdkl5* KO=1.710, SEM=0.05257). **e** MA plot of ELAVLs binding sites in iCLIP experiment shows clear distribution of increased intronic regions (blue) and decreased 3’UTR regions (pink). Each dot represents a single iCLIP binding peak. Dark blue dots and dark pink dots represent significantly changed binding sites. Significance criteria DESeq2 adj.p-value<0.05. n=5 WT mouse brains and 5 *Cdkl5* KO mouse brains. **f** Pie chart distribution of number of RNA features bound by ELAVLs in iCLIP collec-tively in WT and KO mice. Data show preferential binding to 3’UTRs (42%) and introns (44%). **g** Binding motif enrichment analysis shows preferential binding to T-rich (U in RNA) regions as expected for ELAVLs. **h** Comparison of ELAVL iCLIP total intensity (peak count) for mRNA regions show significant reduction in 3’UTR binding and increase in intron binding in *Cdkl5* KOs. n=5 each, WT and KO mouse brains (two-way ANOVA followed by Šídák’s multiple comparisons test, WT (Introns)=20.18, SEM=0.7349; *Cdkl5* KO (Introns)=28.82, SEM=1.823. WT (3’ UTR)=68.65, SEM=3.070; *Cdkl5* KO (3’ UTR)=59.03, SEM=4.085). **i** Representative iCLIP peaks of ELAVLs binding to selected regions in 3’UTR sequence in Camk2a. Individual iCLIP tracks represent each of 5 mouse brains for WT (black) and 5 mouse brains for *Cdkl5* KO (blue). Plus and minus signs indicate the genomic strand. X axis is the genomic location of the iCLIP peak in the mouse GRCm38 genome. **j** mRNA coverage panel shows bulk mRNAseq reads of the same region.

### nELAVL binding to target mRNA 3’UTRs is decreased in *Cdkl5* KO mouse brains

Mammalian ELAVLs reside in both the nucleus and cytoplasm, mediating key functions such as splicing and polyadenylation in the nucleus and enhancing translation in the cytoplasm. To gain insight on how ELAVL function is regulated by CDKL5, we studied ELAVLs’ binding to their target mRNA. Previous studies have identified numerous ELAVL target mRNAs in mouse brain ^52^, human neurons ^51^, as well as in HeLa ^59^ and HEK293 cell lines ^60^. To assess if nELAVL binding to RNA is affected by phosphorylation *in vivo,* we used the improved crosslinking and immunoprecipitation (iCLIP) ^61^ approach with a pan-ELAVL antibody (SC-5261) known to immunoprecipitate ELAVL proteins. This method allowed us to assess transcriptome-wide crosslinking of nELAVLs in *Cdkl5* WT and KO mouse brains at P10, an age of maximal nELAVL phosphorylation **(Fig. 2l)**. We found that the mRNAs bound by ELAVLs included genes with a wide range of functions for neuronal differentiation and synaptogenesis **(Fig. 3e and Supplementary Data 1)**, agreeing with previous findings ^51, 52^. Forty-two percent of the identified iCLIP binding sites were on 3’UTRs, and 44% were on introns (**Fig. 3f and Supplementary Data 1)** and the most enriched binding motifs in iCLIP were T-rich (U in RNA): “TTTTT”, “TTTATT” **(Fig. 3g)**, in agreement with prior data ^51, 52^. We identified 2,123 significantly differentially ELAVL-bound iCLIP binding sites (DESeq2 p-adjusted < 0.05), with 1,626 (76%) of these reduced in *Cdkl5* KOs **(Supplementary Data 1)**. When we plotted the mean normalized counts for the iCLIP sites (indicating the quantity of bound RNA) vs shrunk log2 fold change of binding between WTs and KOs, and incorporated the types of RNA binding site, we observed an opposing trend between intronic and 3’UTR iCLIP binding sites: 3’UTR sites (pink) showed a clear decrease in binding in *Cdkl5* KOs whereas introns (blue) were increased **(Fig. 3e).** When we compared all peak binding intensities for each mouse, we found that 3’ UTR binding was reduced by 1.16 folds and intron binding was increased by 1.43 folds in *Cdkl5* KOs compared to WT mice **(Fig. 3h)**, indicating a global reduction in 3’UTR binding. To characterize total mRNA levels and compare RNA binding data with total RNA levels, we performed bulk mRNA sequencing (RNAseq) experiments on the same brains. CDKL5 mRNA was increased in *Cdkl5* Kos (despite no CDKL5 protein is expressed) and only 11 additional mRNAs were differentially expressed (adj.p-value<0.05) **(Supplementary Fig. 3i and Supplementary Data 1)**, indicating that the iCLIP differences are not due to altered mRNA expression. Representative iCLIP 3’UTR binding sites of ELAVL on calcium/calmodulin dependent protein kinase II alpha (Camk2a) **(Fig. 3i)** with mRNA sequence reads for Camk2a shown below **(Fig. 3j)** demonstrate the specific reduction in ELAVL binding on mRNA without any differences in total mRNA levels. These findings demonstrate that the altered subcellular localization of nELAVLs in *Cdkl5* KO mice is associated with reduced binding to target mRNA 3’UTRs and enhanced binding to intronic regions, which may lead to changes of ELAVL function in these compartments.

### Nuclear roles of nELAVLs are not altered in *Cdkl5* KO mice

We next sought to determine if nELAVLs’ altered localization and mRNA binding affect their nuclear or cytoplasmic functions. Drosophila ELAV and mammalian ELAVLs are known to regulate alternative splicing (AS) and to promote the use of distal alternative polyadenylation (APA) sites in nucleus ^52, 62, 63, 64, 65^. To test if nuclear functions of nELAVLs are affected, we first used Whippet analysis ^66^ to assess changes in RNA splicing events in *Cdkl5* KOs. Our analysis revealed 417 altered events that pass the criteria of probability >0.9, |deltaPsi|> 0.1 and CI width < 0.2, however only 3 nodes had a nearby nELAVL binding site that was changed in *Cdkl5* KOs **(Supplementary Data 3)**, indicating that differential iCLIP binding in introns did not lead to widespread AS. We next carried out APA analysis using 3’RNAseq in *Cdkl5* KO mice at P10. We did not observe any significant changes in APA (adj.p-value<0.05 and change in usage > 10%) (**Supplementary Fig. 3j and Supplementary Data 3**), nor did we observe any trend towards preferred use of distal polyadenylation sites **(Supplementary Fig. 3k)**. These data indicate that an increased nuclear/cytoplasmic nELAVL ratio does not lead to changes in nuclear functions.

### CDKL5 activity regulates new protein synthesis in neurons

ELAV proteins are known to enhance translation of their target mRNAs in the cytoplasm ^67,68^. Since nELAVL cytoplasmic localization was reduced in *Cdkl5* KOs, we next investigated if nELAVL phosphorylation affects mRNA translation. To overcome potential compensatory mechanisms, we used CDKL5 kinase inhibitor CAF-382 to acutely inhibit CDKL5-mediated phosphorylations in rat primary cortical cultures ^26^. Incubation with CAF-382 for 4 hours led to a significant reduction of EB2 phospho-Ser222, indicating that CDKL5 was inhibited **(Fig. 4a,b)**. Inhibiting CDKL5 for 4 hours was sufficient to observe a phosphorylation-dependent increase in nuclear/cytoplasmic ratio of GFP-tagged ELAVL3 when transfected in neurons **(Fig. 4c,d)**. Similar to ELAVL3-GFP, we found increased ELAVL4-GFP nuclear/cytoplasmic ratio upon CAF-382 incubation; S130/131A mutant ELAVL4 did not show a further increase in nuclear/cytoplasmic ratio indicating that CAF-382 acts via ELAVL4 phosphorylation **(Supplementary Fig. 4a-d)**. To measure new protein synthesis, we incubated neuronal cultures with puromycin for 10 minutes and immunostained with anti-puromycin antibody, as previously described ^69^. We found that in the cell body and along the neuronal dendrites there was robust new protein synthesis at 9 days *in vitro* (DIV9) **(Fig. 4e-g)**. Pre-incubation with the translation inhibitor anisomycin eliminated all anti-puromycin staining confirming its specificity **(Fig. 4f).** CAF-382 incubation reduced the puromycin labelling in the cell body and dendrites **(Fig. 4e,g)**, indicating reduced global translation levels. These findings suggest that CDKL5 influences mRNA translation in dendrites, potentially through its regulation of nELAVL phosphorylation and subsequent cytoplasmic localization.

**Fig. 4:**
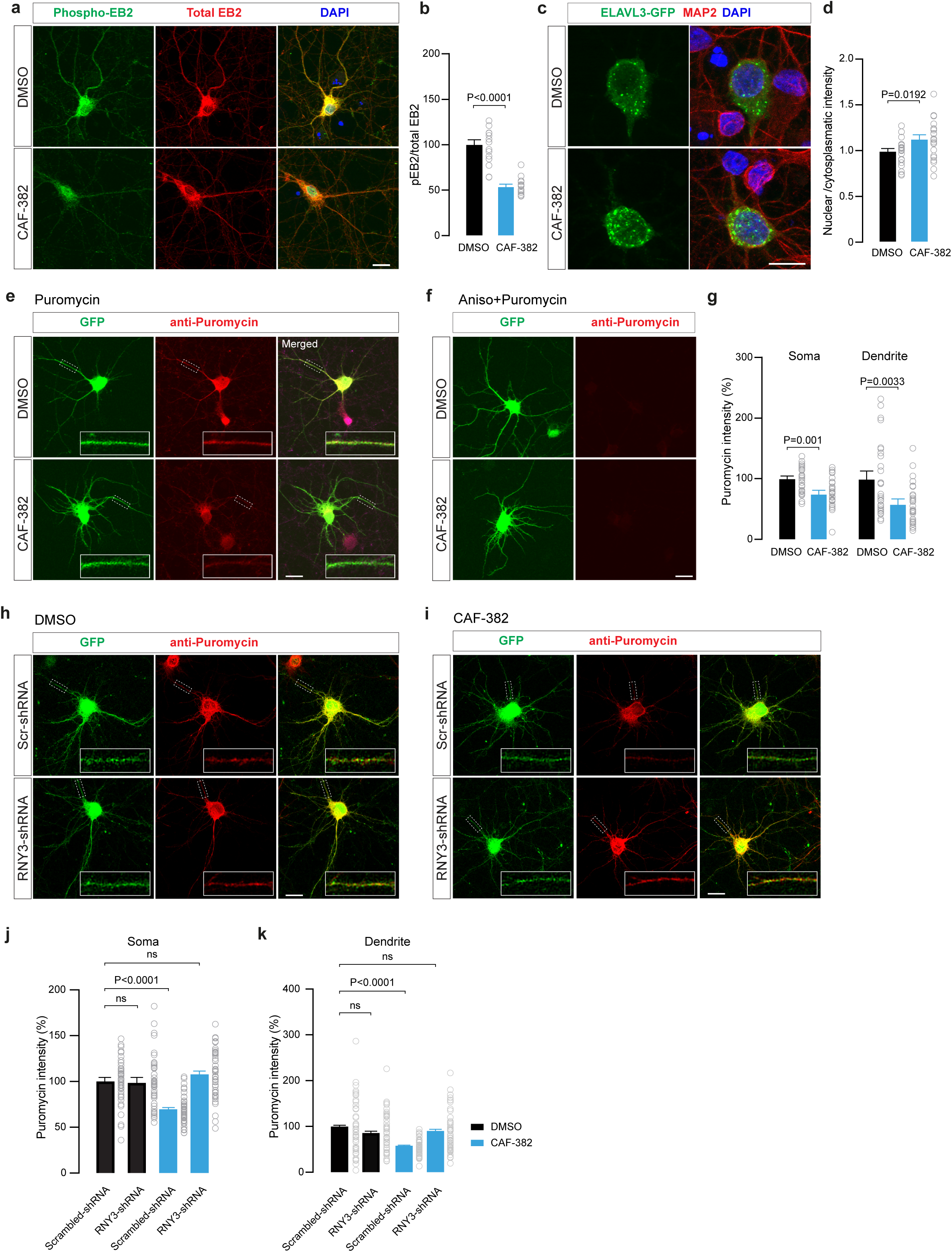
Inhibition of CDKL5 activity regulates ELAVL distribution and translation efficiency. **a** Immunostaining against phospho-EB2 and total EB2 in cortical neurons treated with DMSO or CAF-382 (CDKL5 inhibitor) for 4 hours. Phospho-EB2 is measured in perinuclear region, nuclear staining is non-specific. Scale bar=10 μm. **b** Quantification of the ratio between phospho-EB2 and total EB2 immunofluorescence signals in soma: DMSO, n=15; CAF-382, n=15. (Two-sided unpaired t-test, DMSO=1.000, SEM=5.201; CAF-382 =0.505, SEM=2.445. **c** Distribution of transfected ELAVL3-GFP in neurons treated with DMSO or CAF-382 for 4 hours. Scale bar=20 μm. **d** Graph shows the ratio between nuclear and cytoplasmatic immunofluorescence signals for ELAVL3-GFP. n=21/22 neurons/group. (Two-sided unpaired t-test, Control=100, SEM=3.087; CAF-382=113.9, SEM=4.849). **e** Visualization of newly synthesized proteins with puromycin detected with anti-puromycin antibody in neurons transfected with GFP, after treatment with DMSO or CAF-382 for 4 hours. Scale bar=10 μm. **f** Immunostainings of puromycin labelled proteins (anti-puromycin) in GFP-transfected neurons pre-treated with anisomycin (negative control). Scale bar=10 μm. **g** Quantification from e. DMSO, n= 29; CAF-382, n=29; Two-sided unpaired t-test, Control (soma)= 100, SEM=4.139; CAF-382 (soma)=79.06, SEM=4.369. Control (dendrite)=100, SEM=11.4; CAF-382 (dendrite)=60.17, SEM=6.224). **h, i** Immunostaining against puromycin of DIV9 cortical neurons treated for 4 hours with DMSO (**h**) or CAF-382 (**i**) that have been transduced with scrambled shRNA or RNY3-shRNA for 7 days. Scale bar=10 μm. **j, k** Quantification of mean intensity of puromycin signal within soma (**j**) or dendrites (**k**). One-way ANOVA with Dunnett’s multiple comparisons test. n=40/44 neurons/group. DMSO scrambled-shRNA (soma)=100, SEM=3.582; DMSO RNY3-shRNA (soma)=98.81,SEM=4.679; CAF-382 scrambled-shRNA (soma)=67.91, SEM=2.328; CAF-382 RNY3-shRNA (soma)=106.5, SEM=4.038. DMSO scrambled-shRNA (dendrite)=100, SEM=8.893; DMSO RNY3-shRNA (dendrite)=89.56, SEM=6.906; CAF-382 scrambled-shRNA (dendrite)=57.45,SEM=2.646; CAF-382 RNY3-shRNA (dendrite)=92.80, SEM=6.989.

To test if nELAVL function is involved in CDKL5’s effect in new protein synthesis, we manipulated nELAVL’s function via its negative regulator RNY3. Y RNAs are a conserved family of abundant long non-coding RNAs, with a stem-loop folding. Remarkably, the Y RNA RNY3 binds and sponges the majority of nELAVLs in NSC34 cells ^67^ and in human brain tissue ^51^. Our iCLIP data confirms that RNY3 is the most abundant binder of ELAVLs in P10 mouse brain **(Fig. 3e)**. RNY3 is known to compete with target mRNAs for ELAVL binding ^67^. Inhibiting RNY3 by shRNA-mediated knockdown enhances nELAVL’s binding to target mRNA and their translation ^67^. We examined if RNY3 inhibition can rescue the reduced protein synthesis in the presence of CDKL5 inhibitor by infecting primary cortical neurons with Adeno-associated virus (AAV2) expressing RNY3 shRNA or control shRNA **(Fig. 4h-k)**. We found that protein synthesis reduction caused by CAF-382 were rescued with RNY3 shRNA in soma and dendrites.

In short, by inhibiting a known inhibitor of nELAVL proteins, RNY3, we provided additional evidence supporting our conclusion that nELAVL mediates CDKL5’s positive effect on new protein synthesis. Overall, our findings demonstrate that CDKL5-mediated phosphorylation of nELAVLs during neuronal development promotes the cytoplasmic localization of nELAVLs, where these proteins promote protein synthesis from their bound mRNAs.

### ELAVL2/3/4 phosphorylation is essential for mouse viability

We have identified a novel phosphoregulatory mechanism that affects nELAVL protein function and mRNA translation in neurons. To test if these phosphorylations play a critical role *in vivo*, we generated an *Elavl2/3/4* phosphomutant mouse model using CRISPR/Cas9 approach. We converted both serines at the conserved RPS*S* site to alanines and generated *Elavl2* S118/119A; *Elavl3* S118/119A; *Elavl4* S130/131A mutant mice **(Fig. 5a)**. Each nELAVL was mutated individually and crossed to obtain heterozygous (Het) or homozygous (Hom) phosphomutants. Individual Hom *Elavl2,* Hom *Elavl3* or Hom *Elavl4* phosphomutant (SA) mice were each viable and fertile. When we evaluated double phosphomutants we noticed that double Hom *Elavl2* and *Elavl3* (*Elavl2/3* SA) mice had runty appearance when compared to their littermates and they weighed significantly less than wild-type C57Bl6 controls at P16 (p-value < 0.001, student’s t-test) (**Fig. 5b,c).** Mainly, *Elavl2/3* SA mice did not survive past P20 and due to their runty appearance, most were collected prior to reaching this age. Double Hom *Elavl2/4* mice were viable and fertile and double Hom *Elavl3/4* SA mice were not systematically generated. No triple Hom *Elavl2/3/4* SA mice survived beyond neonatal stage. These findings show that while single Hom nELAVL phosphomutants and dual Hom *Elavl2* and *Elavl4* phosphomutants are tolerated individually, dual Hom *Elavl2* and *Elavl3* is not tolerated. These indicate a more critical role for ELAVL3 than other nELAVLs in agreement with it being the predominant ELAVL3 isoform in brain. These findings show that nELAVL phosphorylations, collectively, are essential.

**Fig. 5:**
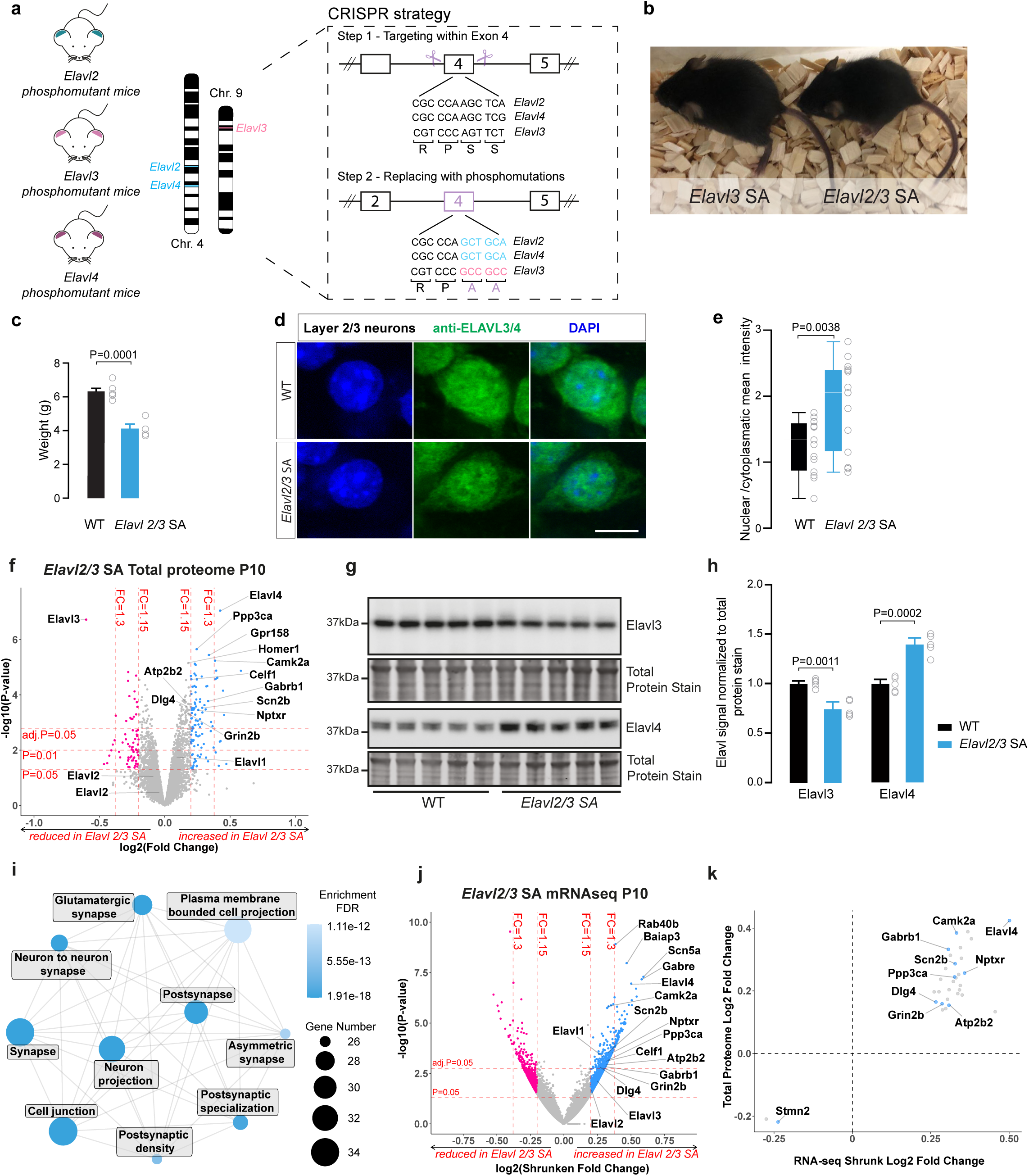
nELAVL phosphorylation is essential for mouse survival, dual *Elavl2/3* phosphomutant mice has compensatory gene regulation. **a** Gene-targeting strategy for generating nElavl (*Elavl*2/3/4) phosphomutant mice. **b** Representative images of P15 homozygous *Elavl*2/3 phosphomutant mice (*Elavl*2/3 SA) and a littermate where only *Elavl3* is homozygous phosphomutant. **c** Body weight in grams at P16. n=4-6 mice/group (Two-sided unpaired t-test, WT=6.322, SEM=0.1858; *Elavl 2/3* SA=4.125, SEM=0.2720). **d** High-magnification view of the layer 2/3 pyramidal neurons from WT and *Elavl2/3* phosphomutant mice at P15 immunostained with an anti-ELAVL3/4. The scale bar is 10μm. **e** Quantification of the nuclear/cytoplasmic mean immunofluorescence ratio shows increase in *Elavl*2/3 phosphomutant mice. n=15 neurons/group used (Two-sided unpaired t-test, WT=1.259, SEM=0.1033; *Elavl 2/3* SA=1.891, SEM=0.1716). **f** Volcano plot of P10 brain proteome comparing *Elavl*2/3 SA and WT using label-free data-independent acquisition (DIA) approach. Each dot represents a different protein. Dots in pink and blue are significantly down- or upregulated, respectively. Criteria for significance are *limma* p-value<0.05 and log2 fold change (lfc)>0.2 = fold change>1.15. n=5 mouse brains for each group. **g** Western blot analysis showing increase of ELAVL3 and decrease of ELAVL4 protein levels at P10 *Elavl*2/3 SA mouse brains. Total protein stain was used as a loading control. n=5 animals per group. **h** Quantification of the Western blots from (**g**). Y-axis is total levels of ELAVL3 and 4 normalised to total protein stain and presented in arbitary units (A.u.). (Two-sided Welch’s t-test, WT (Elavl 3)=1, SEM=0.02317; *Elavl 2/3* SA (Elavl 3)=0.6977, SEM=0.04571. WT (Elavl 4)=1, SEM=0.04361; *Elavl 2/3* SA (Elavl 4)=1.524, SEM=0.06001). **i** Gene ontology analysis using ShinyGO 0.82 for cellular component of significantly upregulated (*limma adj.*p-value<0.05) proteins in P10 Hom *Elavl*2/3 SA mouse brains. Shade of blue indicates the enrichment FDR and the size of the circle corresponds to the number of genes in each node. **j** Volcano plot of P10 bulk mRNA-sequencing comparing *Elavl*2/3 SA and WT. Each dot represents a different gene. Dots in pink and blue are significantly down- or upregulated, respectively. Criteria for significance are DESeq2 p-value<0.05 and log2 shrunken fold change (lfc)>0.2 = fold change>1.15. n=5 animals per group. **k** Quadrant plot showing the correlation of significantly changed proteins (*limma adj.*p-value<0.05) and significantly changed mRNAs (DESeq2 adj.p-value<0.00) in P10 *Elavl*2/3 SA mouse brains. Blue data points indicate overlapping genes where significantly altered binding was observed in *Cdkl5* KO iCLIP analysis (Fig. 3); these mRNAs had reduced 3’UTR binding in *Cdkl5* KO.

Given the lack of respective phosphorylations for Hom phosphomutant mice, we hypothesized that ELAVL subcellular localizations could be more robustly affected in Hom *Elavl2/3* SA mice. Immunostaining for endogenous ELAVL3/4 and ELAVL2 showed increased nuclear localization in layer 2/3 pyramidal neurons in Hom *Elavl2/3* SA (**Fig. 5d,e and Supplementary Fig. 5a,b)**. In addition, hippocampal dentate granule cells, which exclusively express ELAVL3 ^50^, showed a robust increase in nuclear localization in Hom *Elavl2/3* SA mice **(Supplementary Fig. 5c,d)**. These findings confirm the link between nELAVL phosphorylation and their localization *in vivo*. Given ELAVL2 and 3’s robust nuclear enrichment in *Elavl2/3* SA mice we tested if the APA is affected by analysing Hom *Elavl2/3* SA using 3’RNAseq. Similar to *Cdkl5* KOs, we did not observe any significant changes (**Supplementary Fig. 5e and Supplementary Data 3**), showing a lack of effect on ELAVL’s nuclear function.

The severe viability and weight phenotypes observed in homozygous *Elavl2/3* phosphomutant mice highlight the critical physiological importance of these phosphorylation sites. *Cdkl5* KO mouse models do not have any overt phenotypes, yet we know that other kinases, such as CDKL2, can phosphorylate CDKL5 substrates ^55^. Since phosphomutant mice represent a complete loss of the phosphorylation these mouse models can present more severe phenotypes than *Cdkl5* KO mice. Unlike previously reported phosphomutant mouse models of CDKL5 substrates ^27, 70^, nELAVL phosphomutants exhibit severe viability defects. Combined with the evolutionary conservation of ELAVL phosphorylation in Drosophila, these findings strongly suggest that nELAVL phosphorylation represents a critical function of CDKL5.

### *Elavl2/3* dual phosphomutant mice have reduced ELAVL3 levels and trigger altered gene regulation

To study the effects of reduced nELAVL phosphorylations in more detail, we first examined global protein levels in brain. We compared whole-hemisphere brain lysates of P10 Hom *Elavl2/3* phosphomutants to C57Bl6 control mice using label-free data-independent acquisition (DIA) mass spectrometry **(Supplementary Data 4).** The proteome data showed a prominent reduction in ELAVL3 and an increase in ELAVL4 protein levels **(Fig. 5f)**. We confirmed the ELAVL3 reduction and ELAVL4 increase using Western blots. The ELAVL3/4 antibody (Abcam ab184267) showed a reduction at the ELAVL3 molecular weight of 38 kDa, indicating that the antibody predominantly detects ELAVL3 potentially due to its higher abundance when compared to other family members ^52^ and an ELAVL4-specific antibody detected the increase in ELAVL4 levels **(Fig. 5g,h),** confirming mass spectrometry. These findings suggest that loss of ELAVL3’s phosphorylation leads to reduction in its total protein levels, potentially due to protein degradation. In addition, a robust upregulation of ELAVL4 indicates compensatory mechanisms for nELAVL function.

To describe protein level changes in *Elavl2/3* SA mice, we next performed gene ontology (GO) analysis ^71^ for the significantly changed proteome at P10 (adj.p-value<0.05). While the list of significantly decreased proteins did not return any enriched terms, the upregulated proteins (e.g. Camk2a, Grin2b, Gria1) showed robust cellular component (CC) enrichment for postsynaptic density, synapse and specifically glutamatergic synapse **(Fig. 5i and Supplementary Data 4)**. To test if protein changes could be caused by transcriptomics regulations, we next analyzed bulk mRNA sequencing in the same brains. We found that the mRNA level of ELAVL4 was significantly increased (adj.p-value<0.05) **(Fig. 5j and Supplementary Data 4)**, indicating that the increased ELAVL4 protein level is due to an upregulation of *Elavl4* gene expression. We then overlapped our proteomics and transcriptomics data and found a number of common genes/proteins that were significantly increased at both mRNA and protein levels (31 up- and 2 downregulated; **Fig. 5k**). Performing the same GO analysis for the common significantly upregulated genes/proteins returned similar enrichment terms, including glutamatergic synapse **(Supplementary Data 4)**. We next compared these upregulated genes with ELAVL iCLIP data and found that 9 mRNAs which showed reduced ELAVL binding at 3’UTR had compensatory upregulation at transcript and protein level in *Elavl2/3* SA mice, including synaptic proteins Camk2a, Grin2b/GluN2b and Dlg4/Psd95 **(Fig. 5k, blue dots)**. These data show that in the absence of ELAVL2/3 phosphorylation, ELAVL4 upregulation and simultaneous gene expression changes to enable synaptic protein expression, compensating for the reduction in ELAVL2/3 function. Taken together, our results uncover increased abundance of synaptic proteins and their mRNAs in the absence of ELAVL2/3 phosphorylation, which might be an adaptive response to a decrease in their mRNA translation, as a subset of these were also reduced in our 3’UTR iCLIP data.

Overall, analysis of the novel *Elavl2*, *Elavl3* and *Elavl4* phosphomutant mouse models reveals a critical, dose-dependent requirement for these phosphorylation events, and suggests that compensatory mechanisms can be engaged when they are highly reduced.

### Heterozygous *Elavl2/3/4* phosphomutant mice have impaired visual responses

We next investigated the effects of reduced CDKL5-mediated phosphorylation on circuit function. We used triple Het *Elavl2/3/4* SA mice, to model reduced phosphorylation, since *Elavl2/3* SA mice were runty and often did not reach adulthood. We selected visual cortex function as prior work indicated CDKL5 role in visual function in mice ^53, 72^. We characterized the response properties of neurons in the primary visual cortex (V1) of 5 awake, head-restrained Het *Elavl2/3/4* SA mice and 6 WT mice using Neuropixels probes (**Fig. 6a,b**) ^69^. The spontaneous firing rate (FR) in the absence of visual stimulation was similar between the two groups (P = 0.4, Mann–Whitney U test, **Fig. 6c**). To identify potential differences in oscillatory patterns and frequency-specific power between the two genetic groups, we compared the local field potential (LFP) activity for WT and Het animals. We observed consistently lower power across each of the examined frequency bands for the Het *Elavl2/3/4* SA group (P genotype: 0.02, P frequency band: 0.006, P interaction: 0.07, 2-way Anova, **Fig. 6d**). These results are reminiscent of reduced visual responses observed in *Cdkl5* KO mice ^53, 70^, indicating that nELAVLs could be downstream of CDKL5 in regulating visual function.

**Fig. 6:**
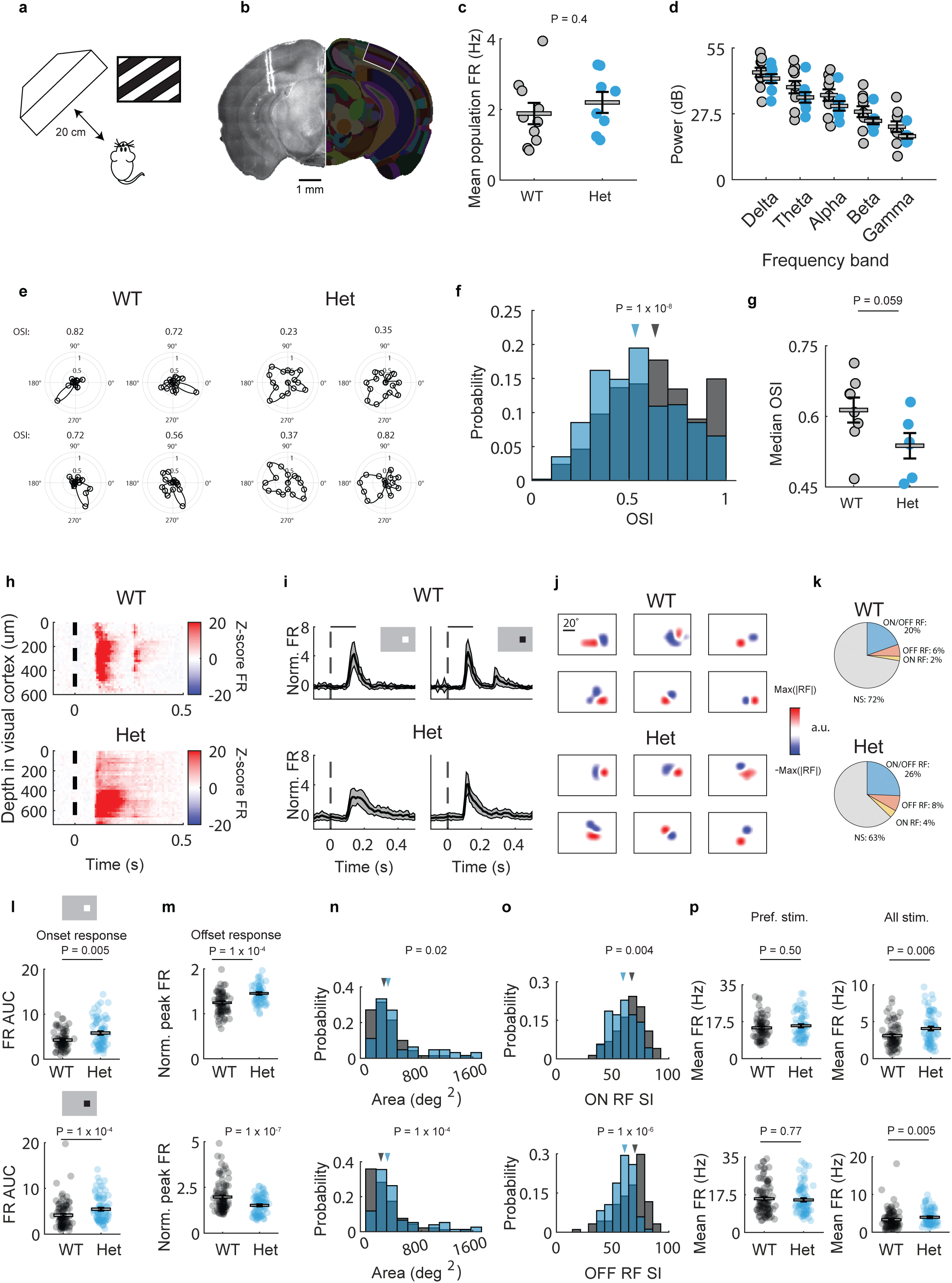
Neuropixels recordings show impaired visual cortex function in *ELAVL2/3/4* triple heterozygous mice. **a** Schematic of in vivo visual stimulus presentation set up. **b** Histological reconstruction of the probe track with DiI coating for an example recording site. Corresponding images from the Allen Mouse Brain Atlas are also present. V1: Primary visual cortex. **c** Quantification of spontaneous population firing rate (FR) for individual recordings. Each dot represents a recording. (WT in black, Het in blue; (Mann–Whitney U = 40, nWT = 10; nHet = 8 recordings, mean ± sem WT: 1.9 ± 0.3, mean ± sem Het: 2.2 ± 0.3, P = 0.4 two-tailed, effect size estimated from rank-biserial correlation, r = 0.250). **d** Power spectrum of LFP analysis on spontaneous activity across different frequency bands: delta (1-4 Hz, Mann–Whitney U = 27, nWT = 10; nHet = 8 recordings, mean ± sem WT: 44.8 ± 1.9, mean ± sem Het: 42.2 ± 1.8, P = 0.27 two-tailed, r = 0.33), theta (4-8Hz, Mann–Whitney U = 26, nWT = 10; nHet = 8 recordings, mean ± sem WT: 38.6 ± 2.6, mean ± sem Het: 34.5 ± 2.2, P = 0.24 two-tailed, r = 0.35), alpha (8-12 Hz, Mann–Whitney U = 23, nWT = 10; nHet = 8 recordings, mean ± sem WT: 35.5 ± 2.4, mean ± sem Het: 20.8 ± 2.0, P = 0.15 two-tailed, r = 0.43), beta (12-30 Hz, Mann–Whitney U = 20, nWT = 10; nHet = 8 recordings, mean ± sem WT: 28.4 ± 2.3, mean ± sem Het: 24.5 ± 1.4, P = 0.08 two-tailed, r = 0.5), gamma (30-100 Hz, Mann–Whitney U = 20, nWT = 10; nHet = 8 recordings, mean ± sem WT: 22.1 ± 2.2, mean ± sem Het: 18.1 ± 1.0, P = 0.08 two-tailed, r = 0.5). **e** Examples polar plots of visual neurons of WT and Het animals. Circular axis spans 360-degree range, distance from the centre of the plot reflects the magnitude of neural responses. Numbers on top of the polar plots indicate orientation selectivity index (OSI) for the example neurons. **f** Distribution of OSI for the two populations. Trian-gles indicate the median of the distribution. (Mann–Whitney U = 98747.50, nWT = 542; nHet = 457 neurons, median ± IQR WT: 0.6 ± 0.34, median ± IQR Het: 0.5 ± 0.33, P = 1×10-8 two-tailed, r = 0.203). **g** Median OSI for individual recordings. (Mann–Whitney U = 9.00, nWT = 8; nHet = 6 recordings, mean ± sem WT: 0.6 ± 0.02, mean ± sem Het: 0.5 ± 0.02, P = 0.059 two-tailed, r = 0.625). **h** Average spiking response to visual stimulus during receptive field mapping across the depth of V1. Neurons are grouped in 20um bins. Dashed black line indicates stimulus onset. Colour bar indicates z-scored FR. **i** Mean population firing rate during white (left) and black (right) patches for corresponding recordings in (**h**). Shaded area indicates standard deviation. **j** Combined ON and OFF RF subfield maps of examples visually tuned neurons. Colour intensity denotes response strength to light increments (ON, red) and light decrements (OFF, blue). **k** Pie chart representing percentage of responsive neurons in the two populations (ON RF: neuron significantly responding to white patches; OFF RF: neurons significantly responding to black patches; ON/OFF RF: neurons signifi-cantly responding to both stimuli type; NS: not significant). **l** Area under the curve (AUC) of onset response (50ms - 250ms after stimulus) during white (top) or black (bottom) patches. Each data point corresponds to a V1 neuron. (For white patch: Mann–Whit-ney U = 1883.50, nWT = 74; nHet = 70 neurons, mean ± sem WT: 4.3 ± 0.2, mean ± sem Het: 5.8 ± 0.4, P = 0.005 two-tailed, r = 0.273; For black patch: Mann–Whitney U = 2765, nWT = 94; nHet = 85 neurons, mean ± sem WT: 4.2 ± 0.3, mean ± sem Het: 5.4 ± 0.3, P = 1×10-4 two-tailed, r = 0.308). **m** Normalised firing rate during offset responses (250ms – 400ms after stimulus). (For white patch: Mann–Whitney U = 2780, nWT = 74; nHet = 70 neurons, mean ± sem WT: 1.3 ± 0.03, mean ± sem Het: 1.5 ± 0.03, P = 1×10-4 two-tailed, r = 0.304; For black patch: Mann–Whitney U = 1338, nWT = 94; nHet = 85 neurons, mean ± sem WT: 2.0 ± 0.08, mean ± sem Het: 1.5 ± 0.05, P = 1×10-7 two-tailed, r = 0.483). **n** Distribution of ON and OFF subfield area. (For white patch: Mann–Whitney U = 1702.50, nWT = 70; nHet = 63 neurons, median ± IQR WT: 309.1 ± 243.6, median ± IQR Het: 362.9 ± 274.3, P = 0.02 two-tailed, r = 0.228; For black patch: Mann–Whitney U = 2428, nWT = 92; nHet = 76 neurons, median ± IQR WT: 270.1 ± 270.2, median ± IQR Het: 255.2 ± 164.6, P = 1×10-4 two-tailed, r = 0.305). **o** Distribution of selectivity indices for white and black patches. (For white patch: Mann–Whitney U = 1871, nWT = 74; nHet = 70 neurons, median ± IQR WT: 67.1 ± 17.5, median ± IQR Het: 59.3 ± 17.6, P = 0.004 two-tailed, r = 0.278; For black patch: Mann–Whitney U = 2368, nWT = 94; nHet = 85 neurons, median ± IQR WT: 69.9 ± 16.0, median ± IQR Het: 60.9 ± 14.5, P = 1×10-6 two-tailed, r = 0.407). **p** Mean FR at the preferred stimulus location (left column) and mean FR across all stimuli locations (right column). For the preferred stimulus location, white patch: Mann–Whit-ney U = 2422.50, nWT = 74; nHet = 70 neurons, mean ± sem WT: 14.6 ± 0.6, mean ± sem Het: 15.6 ± 0.7, P = 0.50 two-tailed, r = 0.065; For black patch: Mann–Whitney U = 3895, nWT = 94; nHet = 85 neurons, mean ± sem WT: 15.7 ± 0.75, mean ± sem Het: 15.1 ± 0.7, P = 0.77 two-tailed, r = 0.025; for all stimuli locations, white path: Mann–Whitney U = 1903.50, nWT = 74; nHet = 70 neurons, mean ± sem WT: 3.1 ± 0.2, mean ± sem Het: 4.1 ± 0.3, P = 0.006 two-tailed, r = 0.265; For black patch: Mann–Whitney U = 3014.50, nWT = 94; nHet = 85 neurons, mean ± sem WT: 3.3 ± 0.3, mean ± sem Het: 3.9 ± 0.2, P = 0.005 two-tailed, r = 0.245).

To assess potential differences in visual response properties between WT and Het animals we recorded neuronal responses to square wave drifting gratings and calculated the orientation selectivity index. While, both wild-type and Het *Elavl2/3/4* SA animals displayed visually evoked responses (**Fig. 6e**), visual inspections of the neuronal responses revealed differences in the degree of selectivity for specific orientations and directions of movement: visual neurons of the Het *Elavl2/3/4* SA mice show a broader distribution of response magnitudes across multiple orientation. Consistent with these results, neurons in the Het *Elavl2/3/4* SA animals displayed lower orientation selectivity than neurons in their WT counterparts when pooling data across animals (median OSI Het: 0.53, median OSI WT: 0.63, P: 1 × 10^−8^, Mann–Whitney U test, **Fig. 6f**). However, differences between the median OSI of the two groups fell short of reaching significance (P: 0.059, Mann–Whitney U test, **Fig. 6g**). Similar results were observed for the direction selectivity index (DSI) **(Supplementary Fig. 6).**

To further characterise visual properties of V1 neurons of Het *Elavl2/3/4* SA mice we mapped the structure of spatial receptive fields (RFs) using sparse noise stimuli. Visually evoked responses were observed along the depth of V1 (**Fig. 6h**) for both groups of animals. However, we observed longer responses evoked by the onset of the stimuli in the triple Het recordings, accompanied by an absence of offset responses (**Fig. 6i**). Despite the differences in the response dynamics of the two groups, we found neurons that exhibited significant spatial RFs in both populations (**Fig. 6j**) and we observed a similar proportion of responsive neurons in both populations (**Fig. 6k**). The differences in neuronal response dynamics in neurons in the Het *Elavl2/3/4* SA animals were characterised by larger onset responses compared to neurons in their WT counterparts (white patches: P: 0.005; black patches: P: 1 × 10^−4^, Mann–Whitney U test, **Fig. 6l**). These higher responses of individual neurons are consistent with the prolonged population firing rates observed in the example of panels **6h** and **6i.** Interestingly, neurons from Het *Elavl2/3/4* SA mice also displayed a lower FR during the offset response to the black patches than neurons from wild-type mice (P: 1 × 10^−7^, **Fig. 6m**). However, because offset responses to white patches merged with the prolonged onset responses of the V1 neurons of Het *Elavl2/3/4* SA animals, the net effect was an overall higher firing rate compared to the neurons from the WT mice (P: 1 × 10^−4^, Mann–Whitney U test, **Fig. 6m**).

Further analysis of RF properties revealed that neurons in the Het *Elavl2/3/4* SA animals showed larger RF areas than WT for both ON and OFF subfields (median ON RF size: WT: 309, Het: 362, P: 0.02; median OFF RF size: WT: 270, Het: 355, P: 1 × 10^−4^, Mann–Whitney U test, **Fig. 6n**) and lower spatial selectivity than those of the wild-type animals (median ON RF SI: WT: 67, Het: 59, P: 0.004; median OFF RF SI: WT: 69, Het: 60, P: 1 × 10^−6^, Mann–

Whitney U test, **Fig. 6o).** The lower selectivity index was not explained by a change in the peak firing rate to the preferred location (white patches: P: 0.50; black patches: P: 0.77, Mann–Whitney U test, **Fig. 6p**), but resulted from an increase in the neuronal responses of Het *Elavl2/3/4* SA animals to all stimuli locations (white patches: P: 0.006; black patches: P: 0.005, Mann–Whitney U test, **Fig. 6p**). Together these results show altered visual response properties of V1 neurons in Het *Elavl2/3/4* SA mice, evidenced by a reduction in orientation and RF selectivity.

Collectively, our findings uncover a novel role for CDKL5 in phosphorylating conserved neuronal ELAVL proteins, thereby promoting their cytoplasmic localization, mRNA binding, and protein synthesis **(Fig. 7)**. This phosphorylation, highly active during development, is crucial for the proper formation of visual circuits, highlighting a novel mechanism in neurodevelopment.

**Fig. 7:**
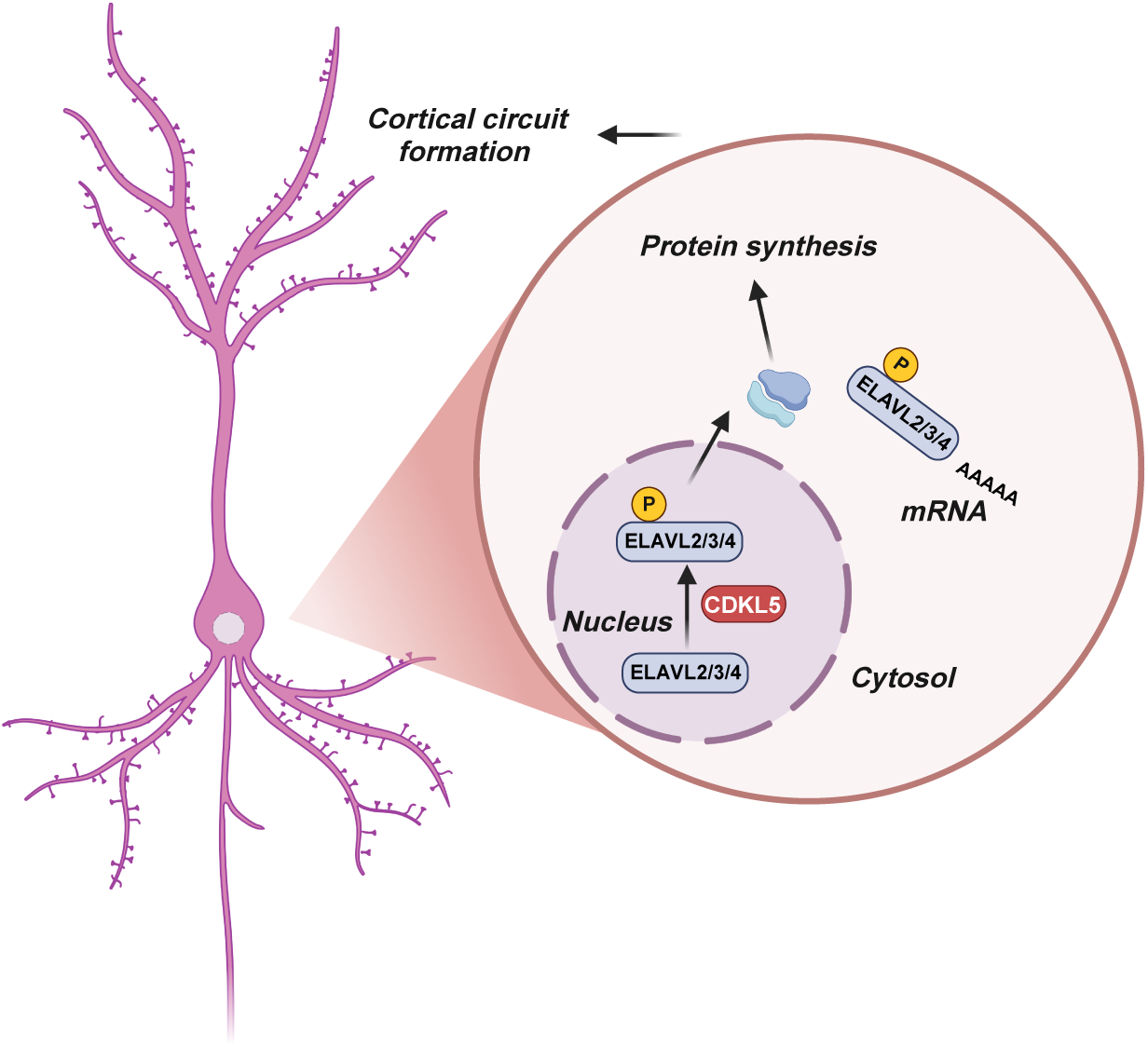
Regulation of neuronal ELAVL proteins by CDKL5. CDKL5-dependent phosphorylation of neuronal ELAVL proteins regulates their shuttling to cytoplasm, facilitating their function in mRNA binding, protein synthesis and development of visual cortex circuits.

## Discussion

Using global phosphoproteomics analysis, we report for the first time a comprehensive list of CDKL5 substrates and their phosphorylation sites, all of which share the CDKL5 consensus motif. The diverse subcellular localizations and known functions of these substrates suggest that CDKL5 is involved in a wide range of cellular processes. To fully understand the functional spectrum of CDKL5, further investigation of its substrates will be needed. CDKL5’s phosphorylation site on nELAVL is conserved between all mammalian neuronal ELAVL proteins as well as in Drosophila homologs ELAV (S242) and FNE (S121) proteins. In Drosophila and mammalian nELAVL proteins serine 119/131 phosphorylation site is located between RRM1 and RRM2. Using differentiated human iPSC-derived neurons from patients and parents we were able to demonstrate that nELAVL phosphorylation is also reduced in human neurons with *Cdkl5* deletion. The remaining nELAVL phosphorylation in mice and humans can be due to other kinases including CDKL2 ^55^. These results suggest a conserved phosphoregulatory mechanism. The developmental time course of p-nELAVL shows a peak in the first two postnatal weeks in mice, in agreement with EB2 and MAP1S phosphorylation time course ^13^ indicating a critical role for nELAVL phosphorylation during neuronal differentiation.

To reveal the role of nELAVL phosphorylation we studied phosphomutant proteins in multiple functional assays *in vitro* and *in vivo*. Our *in vitro* RNA-ELAVL4 binding experiments did not reveal any changes in affinity due to phosphorylation. In support of this, the phosphorylation of ELAVL1 (HuR) S100 (RPSS*) in the interdomain linker region is not expected to affect RNA binding in crystal structure models ^73^. Results using single or dual phoshomutant ELAVL4 overexpression in NSC34 cells indicate a dose dependent effect of phosphorylation at the RPS*S* site on nELAVL cytoplasmic localization. Using two mouse models of ELAVL phosphorylation deficiency, *Cdkl5* KO and Hom *Elavl2/3* SA, we demonstrated that phosphorylation by CDKL5 facilitates nELAVL cytoplasmic localization *in vivo*. Our results demonstrate an increased nuclear/cytoplasmic ratio using antibodies that detect endogenous ELAVL3/4 and ELAVL2. Interestingly, in dentate granule neurons of the hippocampus, which only express ELAVL3 ^52^, we observed a similar increased nuclear/cytoplasmic staining in *Cdkl5* KO and Hom *Elavl2/3* SA mice, showing that ELAVL3 is specifically regulated by CDKL5. ELAVL2 showed a similar shift to nucleus in both mutant mice, albeit more cytoplasmic that ELAVL3/4. It has been demonstrated that the hinge region between RRM2 and RRM3 of ELAVLs harbours signals facilitating both nuclear export and nuclear localization ^32, 74^. Our results are the first indication that the interdomain linker region between RRM1-RRM2 also play a role in cytoplasmic localization.

Phosphorylation may affect nELAVL interaction with export factors which is characteristic for export adaptors, RNA-binding proteins that handover mRNAs to TAP/NXF1 and licence it for nucleocytoplasmic transport ^75^. The molecular mechanisms of phosphorylation’s effect on nuclear/cytoplasmic localization remain to be determined.

We observed a significantly decreased ELAVL binding to 3’UTR of numerous mRNAs and an increase in nELAVL binding to introns, mimicking the endogenous nELAVL mislocalizations in *Cdkl5* KO and *Elavl2/3* SA mice. Increased intron binding to the unprocessed RNA in the nucleus could potentially cause changes in nELAVLs’ function in RNA processing, such as APA. Our extensive analysis using bulk or 3’ RNA sequencing data showed no significant changes in APA or in 3’UTR lengths in *Cdkl5* KO or Hom *Elavl2/3* SA mice, unlike the changes seen previously in the complete absence of ELAVL ^63, 65, 76^. Thus, nuclear functions of nELAVLs do not seem to be affected in our mouse models. ELAVL proteins are also known to regulate RNA stability, which leads to changes in mRNA abundance ^77, 78^. Since we detected few significant differences with minor fold-changes in the RNAseq experiment from *Cdkl5* KOs **(Supplementary Fig. 3i),** we did not further investigate the RNA stability.

Protein synthesis and homeostasis contribute to neurodevelopmental disorders^79^. A major role of ELAVL proteins is to enhance translation efficiency of target mRNAs, a function which relies on their mRNA 3’UTR binding in the cytoplasm ^67, 68^. Our data reveal significant reductions in 3’UTR binding by ELAVL in *Cdkl5* KO. Surprisingly, we do not observe differences in total protein levels that correlate with the mRNA bound by nELAVLs in *Cdkl5* KO mice as the proteome remains largely unchanged. Total protein levels may be compensated by changes in mRNA translation or protein turnover pathways. Alternatively, analysis of protein levels in brain hemispheres may have masked differences which could have been detectable in individual brain regions, protein levels of other cell types could have masked neuron-specific alterations or our mass spectrometry data could be underpowered. To address CDKL5’s role in protein synthesis more effectively, we examined the acute loss of nELAVL phosphorylations in primary cortical neurons using a specific CDKL5 inhibitor ^26^. We found that CDKL5 activity is necessary for efficient protein synthesis. By demonstrating an epistatic interaction between CDKL5 inhibition and RNY3 knockdown, we further determined that CDKL5 acts via nELAVLs to facilitate protein synthesis. Notably, knockdown of RNY3 does not affect puromycin incorporation in control neurons; this could be because nELAVL binding to target mRNAs may not be a rate-limiting step under baseline conditions of neuronal development, thus enhancing such binding does not increase protein synthesis further. In contrast to *Cdkl5* KOs, in Hom *Elavl2/3* phosphomutant mice we observed a surprising upregulation of ELAVL4 mRNA and protein, in parallel to an upregulation of multiple mRNAs and proteins for excitatory synapse components. These compensatory mechanisms could lead to dysregulation of neuronal synaptogenesis and could represent a molecular state caused by loss of nELAVL phosphorylation, which could be present in CDD patients. Future studies would be needed to determine how RNA binding, localization or translational control are regulated by CDKL5 and nELAVLs in human neurons.

Generating novel mouse models is essential in determining the functional roles of proteins and phosphorylations in organisms. The *Elavl3* full knockout (KO) mouse model is viable and fertile but exhibits spontaneous epilepsy ^52^. In contrast, double *Elavl2/3* KO mice die shortly after birth, highlighting the essential roles of these genes in brain development ^52^. The triple CRISPR -mutated *Elavl2/3/4* phosphomutant mouse models we generated indicated that nELAVL are critical substrates phosphorylation for CDD pathology. To investigate the role of nELAVL phosphorylations in circuit development *in vivo*, we used the triple heterozygous *Elavl2/3/4* phosphomutant mice. Our detailed analysis of visual response properties in these mice revealed impaired visual responses, including reduced selectivity for orientation and receptive field preference, supporting the importance of nELAVLs in visual system development. CDD patients often present with cortical visual impairment (CVI) ^80^ and diminished visual electrophysiological responses in the cortex ^81^, while *Cdkl5* KO mice also show reduced visual response magnitude ^53, 72^, mirroring our findings. A visual discrimination task could be used in the future in order to measure visual function in Elavl2/3/4 heterozygous mice. These results suggest that nELAVL phosphorylation by CDKL5 is a crucial aspect of CDKL5 function, potentially explaining the visual defects observed in CDD pathology.

Collectively our data reveal a key phosphorylation mechanism for a neurodevelopmental disorder associated kinase, CDKL5, in controlling protein synthesis and visual cortical circuitry by phosphorylating the ubiquitous RNA binding proteins neuronal ELAVLs.

## METHODS

### Animals

The mice were bred and handled according to the regulation of the animal (Scientific Procedures) Act 1986 of the United Kingdom, and approved by the Francis Crick Institute Animal Welfare and Ethical Review Body (AWERB). The mice were housed and maintained on a 12 hour light/dark cycle and provided with food and water *ad libidum*. Each mouse strain was backcrossed into C57BL/6J genetic background.

*Cdkl5 full body knockout mice.* These mice were a generous gift from Cornelius Gross at the European Molecular Biology Laboratory in Rome^53^. Heterozygous females and wild-type males were used to maintain *Cdkl5* full KO mouse line and only the brains of male offspring were harvested to ensure *Cdkl5* WT and hemizygous KO littermates.

#### ELAVL2/3/4 phosphomutant mice

Two guide RNAs were used for each mutation. All CRISPR guides were selected to have at least three base-pair mismatches relative to the genome. For one *Elavl2* guide, the closest predicted off-target contained a two-base-pair mismatch and was located within an intron. For all guides except this *Elavl2* guide, the predicted three-base-pair mismatches mapped to different chromosomes. In the case of the *Elavl2* guide for which the three-base-pair mismatches mapped to the same chromosome, these sites would still be expected to segregate independently. Therefore, any potential off-target mutations would be expected to segregate independently.

Mice carrying phosphomutant alleles of *Elavl4* and *Elavl2* were generated concurrently, whereas *Elavl3* phosphomutant lines were generated independently. All lines were initially crossed to the C57BL/6 background to expand colonies, and no line was maintained as homozygous. Importantly, homozygous *Elavl2/4* phosphomutant mice were viable and fertile and exhibited no overt phenotypes, effectively ruling out off-target effects as a cause of reduced body weight. Similarly, homozygous *Elavl3* phosphomutant mice were viable, fertile, and phenotypically normal, further indicating the absence of off-target mutations associated with decreased body weight.

Progeny from the *Elavl2/4* line were subsequently crossed with the *Elavl3* line to generate multiple genotypes, which were then intercrossed. The colony has undergone repeated intercrossing and backcrossing to C57BL/6 mice.

Detailed CRISPR strategy of each mutation is provided below.

### Generation of *Elavl3* phosphomutant mouse model

Two guide RNAs (sgRNAs) were designed using CRISPOR (http://crispor.tefor.net/) to cut in exon 4 (ENMUSE00000285207) of the *Elavl3* gene (Table 1). A 102 bp ssODN (single stranded oligonucleotide) donor molecule was designed to be used with either guide and contained the phosphomutations S118A (AGT>GCC) and S119A (TCT>GCC) mutations, an additional silent mutation for each guide, and 47-49 bp homology arms either side (Table 2). C57BL/6J zygotes were electroporated with 50 µL of Cas9 protein/sgRNA/ssODN (1.2 µM / 6 <u>µ</u>M / 8 <u>µ</u> M) using a Nepa21 electroporator (Sonidel Ltd^TM^).

Preliminary Transnetyx qPCR was performed on mouse ear-snips to identify nine F0 mice positive for the mutation. All founder mice were subjected to 399 bp PCR across the entire ssODN and high-throughput sequencing of the PCR amplicons was performed by the Advanced Sequencing Facility using Illumina MiSeq sequencing 250 bp paired-end reads (Table 3). A single founder mouse with precise integration of the point mutations was identified using sgRNA-1 and a second founder mouse with the same precise mutation was identified using sgRNA-2. The two F0 mice were bred to C57BL/6J which resulted in three and two F1 mice respectively, identified as heterozygous for the intended mutation by MiSeq (Illumina, Inc) amplicon sequencing. Oxford Nanopore Technologies (ONT) sequencing was performed on 3 kb amplicons, generated by PCR, spanning the targeted insertion in one F1 mouse from each line. No aberrant on-target mutations or deletions were found either 1.5 kb upstream or downstream of the insertion site in either.

### Generation of *Elavl2/4* double phosphomutant mouse model

Two guide RNAs (sgRNAs) were designed using CRISPOR to cut in exon 4 (ENMUSE00001307484) of the *Elavl2* gene. A 120 bp ssODN donor molecule was designed to be used with either guide and contained the S118A (AGT>GCT) and S119A (TCA>GCA) mutations, silent mutations for the respective guides, and 57 bp homology arms on either side (Table 2). Two guide RNAs (sgRNAs) were designed using CRISPOR to cut in exon 4 (ENMUSE00001208755) of the *Elavl4* gene. A 120bp ssODN donor molecule was designed to be used with either guide and contained the S130A (AGC>GCT) and S131A (TCG>GCT) mutations, silent mutations for the respective guides, and 53-61 bp homology arms on either side (Table 2).

Both ssODN, Elavl2 sgRNA-2 and Elavl4 sgRNA-2 were electroporated into C57BL/6J zygotes with 50uL of Cas9 protein/(total)sgRNA/(total)ssODN (1.2 <u>µ</u> M / 6uM/8uM) using a Nepa21 electroporator. One F0 mouse was born and genotyped as an Elavl2 negative/Elav4 homozygote by Transnetyx qPCR. A PCR of 384bp over the Elavl2 and Elavl2 340bp over each insertion site was performed on the F0 (Table 3). Subsequent MiSeq (Illumina, Inc) amplicon sequencing analysis revealed a homozygous insertion of the *Elavl4* phospho-mutation, the *Elavl2* mutation was not identified however there was integration of one silent mutation on one allele and on the other allele a spurious 2bp deletion. Zygotes homozygous for the *Elavl4* phospho-mutation were generated by in vitro fertilization and were subsequently electroporated with Cas9 protein/Elavl2 ssODN/Elavl2 sgRNA-2 as previously described. The resulting mice were screened by Transnetyx, and three founder mice were identified positive for the intended *Elavl2* mutation and the previously introduced *Elavl4* mutation. The correct integration of the mutations at both sites was confirmed by MiSeq (Illumina, Inc). amplicon sequencing, and all three founders were bred with C57BL6/J, resulting in three F1 double phosphomutant mice. Oxford Nanopore Technologies (ONT) sequencing was performed on 3kb amplicons, generated by PCR, spanning the targeted insertions on one F1 mouse. No aberrant on-target mutations or deletions were found 1.5kb upstream or downstream of either insertion ^82^.

**Methods table 1.**
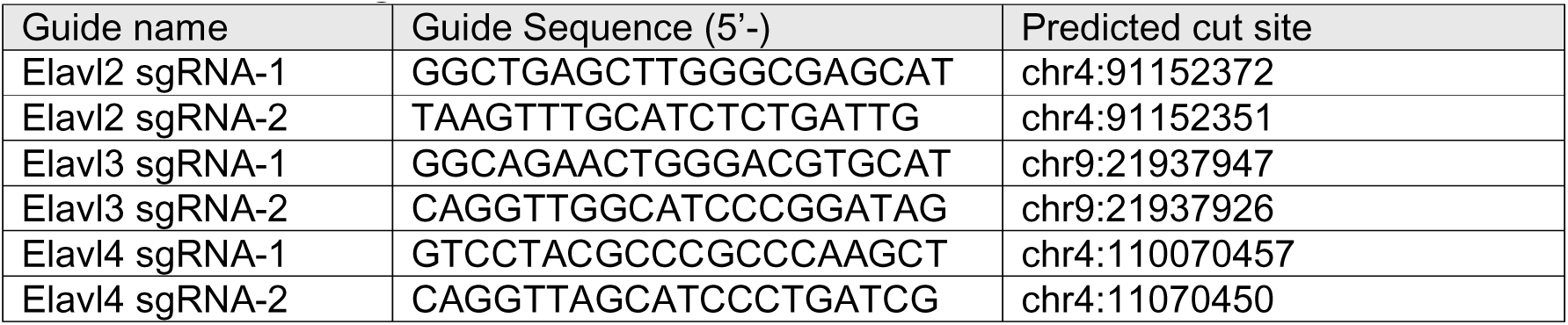
sgRNA.

**Methods table 2.**
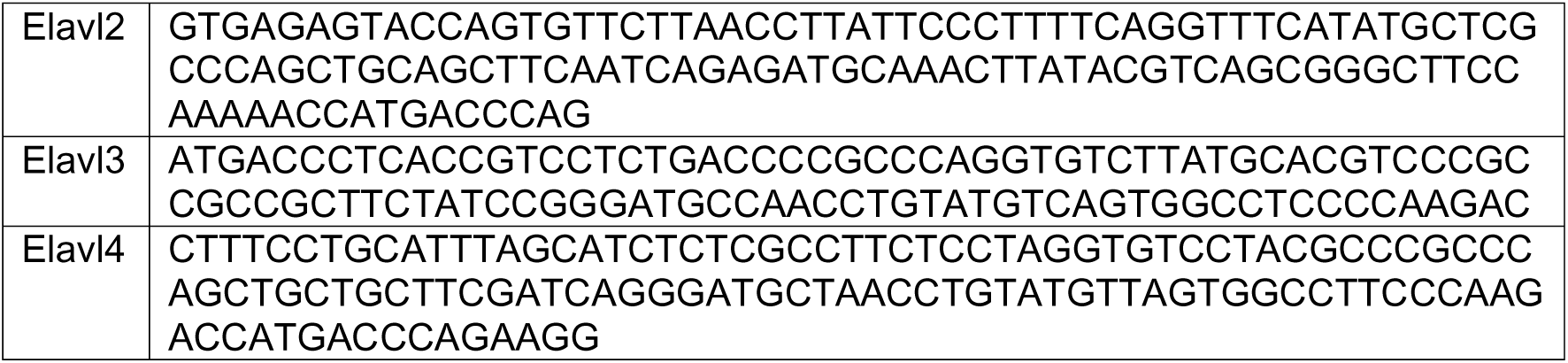
ssODN Sequences.

**Methods table 3.**
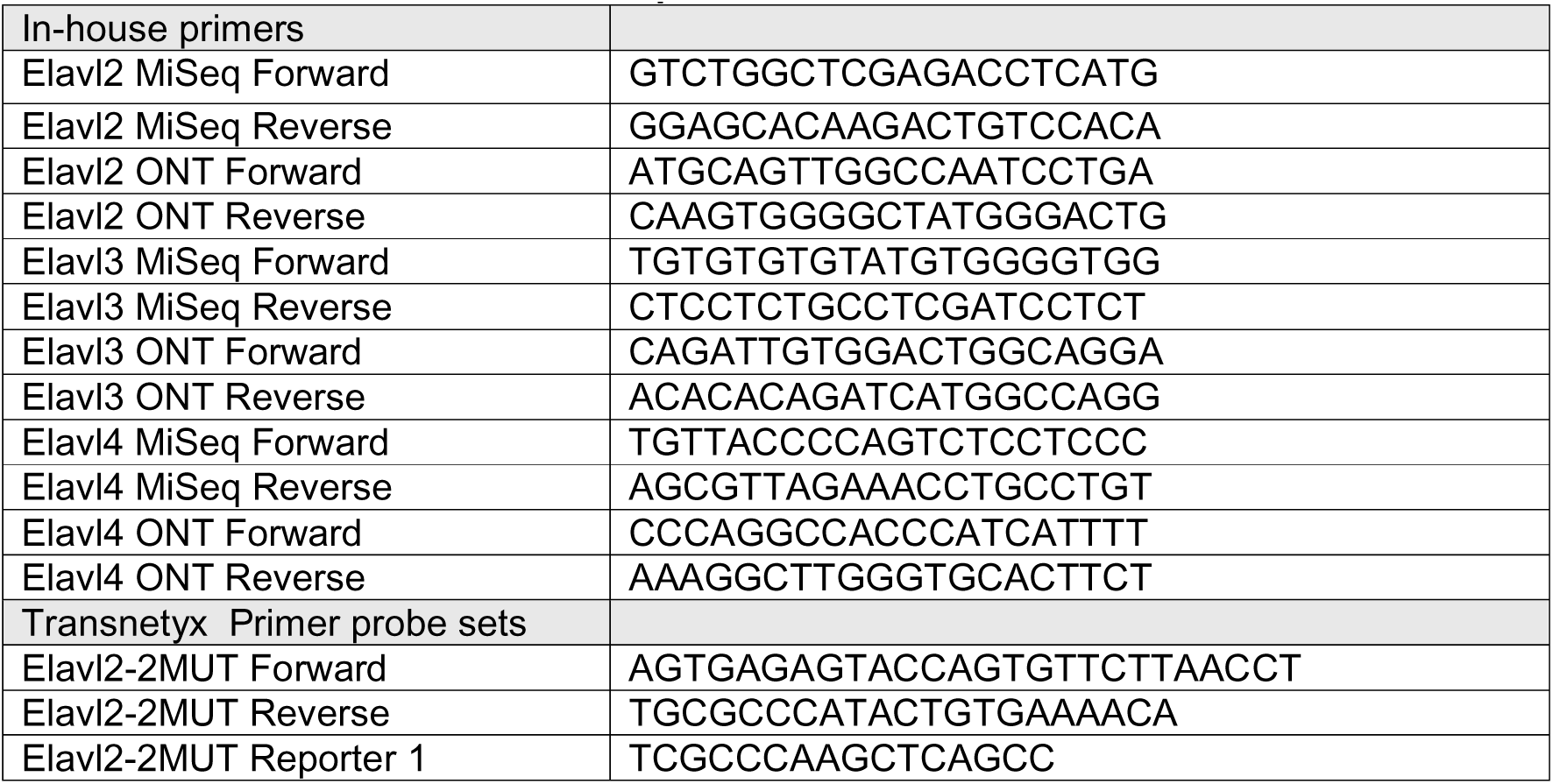
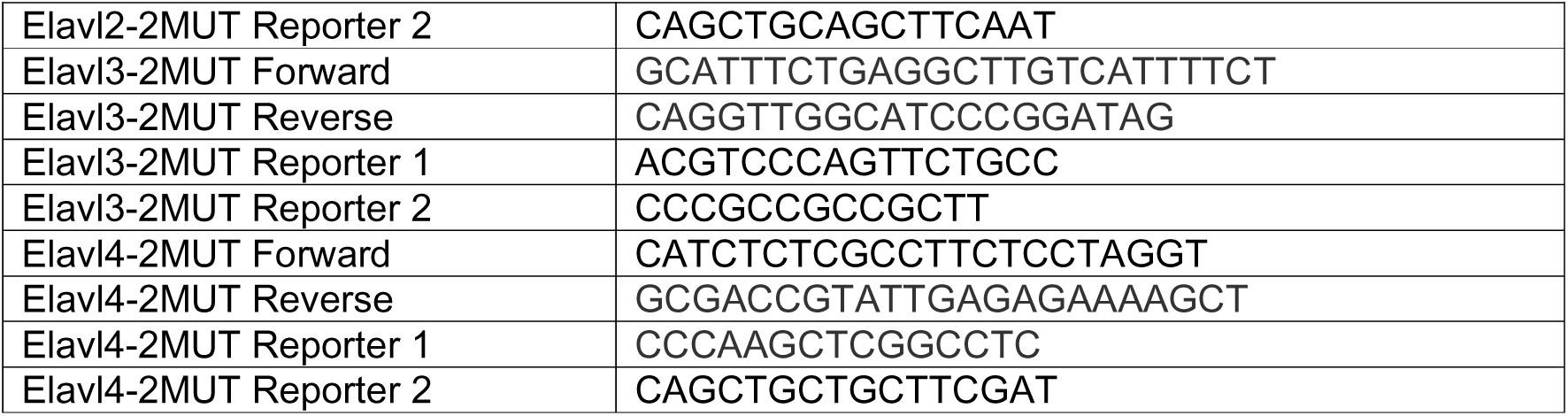
Primer/Probe Sequences.

### Histology

*Western blots*. Fresh brains were taken promptly from mice after cervical dislocation and decapitation. When desired, brain regions were dissected and tissue was flash frozen in liquid nitrogen (LN2).

*Brain tissue section staining.* Mice were anesthetized and transcardial perfused with ice-cold PBS followed by 4% ice-cold paraformaldehyde (PFA). Brains were dissected out and post-fixed in 4% PFA overnight and stored in PBS-azide at 4°C. Brains from WT and Elavl2/3 SA mice were embedded in 4% agarose and sectioned into 50 µm coronal sections using a vibratome (Leica VT1000 S). Brain sections were collected in 0.02% PBS-azide and stored at 4 °C.

### Cell cultures

*HEK293T and NSC34 cells*. The cells were grown in Dulbecco’s modified Eagle medium (DMEM) supplemented with FBS and penicillin/streptomycin at 37°C with 5% CO_2_ for maximum 25 passages. Each passage consisted in dissociating the cells using 0.25% trypsin-EDTA for 3 minutes at 37°C and adding 1:10 of the cells into a new T75 flask. This step was repeated every 3-4 days.

*Patient-derived neuron lysates.* Induced pluripotent stem cells (iPSCs)-derived neuron lysates were a generous gift from Dr. Allyson Muotri at the University of California, San Diego, reported in ^56^. Briefly, iPSC media was replaced with DMEM/F12 (Corning Cellgro) containing 1X HEPES, 1X penicillin-streptomycin, 1X Glutamax (Life Technologies) and 1X N2 NeuroPlex (Gemini Bio-products), supplemented with 1 mM dorsomorphin (Tocris) and 10 mM SB431542 (StemGent). Next, colonies were kept in suspension for 7 days to form embryoid bodies, which were plated onto matrigel-coated plates and cultured using DMEM/ F12 with 1X HEPES, 1X penicillin-streptomycin, GlutaMAX, 0.5X N2 NeuroPlex and 1X Gem21 NeuroPlex (Gemini Bio-products), supplemented with 20 ng/mL bFGF (Life Technologies). Neural rosettes were manually collected, gently dissociated with accutase (Stem Cell Technologies), and the neural progenitor cells (NPCs) were replated on poly-L-ornithine/laminin-coated plates. Next, the NPCs were differentiated into neurones by adding 5 μM ROCK inhibitor (Tocris) for 48 hours and bFGF withdraw for 6 weeks. The cells were then collected, pelleted and the pellet was frozen in LN2. The pellet was lysed in 1X sample buffer (Invitrogen) containing 0.1 M dithiothreitol (DTT), sonicated briefly twice and denatured at 70°C for 10 minutes. The samples were finally centrifuged at 13,300 rpm for 10 minutes and a Western blot was performed following the protocol described below (Western Blot). CDD Patient 1 is a female patient with the R59X mutation and the control parent is her mother. CDD Patient 2 is a male patient with c.404-1G>A and p.D135_F154del mutations and the control parent is his father ^56^.

### DNA constructs

pLVExp-CMV-EGFP-rELAVL4 WT plasmid was purchased from VectorBuilder. rELAVL4 phosphomutant (SA) plasmid was generating by introducing the mutation S131A using site-directed mutagenesis. pRK5-HA-CDKL5_1-352_ (kinase domain) WT and kinase dead (KD) vectors, as well as the pFastBacHT-His6-CDKL5_1–352_ vector, were already described ^13^. Full-length mELAVL4 in a pGEX3X vector (generously donated by Prof. Toshinobu Fujiwara from Kindai University) was cloned into a pET47b-His6 vector. mELAVL4 was then truncated using inverse PCR and only the first two domains from amino acid 14 to 215 were kept (pET47b-His6-mELAVL414-215 or RRM1/2 mELAVL4).

### ELAVL2/3/4 S119/131 phosphospecific antibody

Rabbit polyclonal ELAVL2/3/4 phosphospecific antibodies were raised against the following phosphorylated (*) peptides by Covalab: IKVSYARPSS*ASIR. The peptide sequence is common to ELAVL2, ELAVL3 and ELAVL4 but does not match ELAVL1. New Zealand White rabbits were injected with the phosphorylated peptide. Final bleeds of immunized animals were purified with affinity purification by Covalab. In short, the immune serum was loaded onto a column with the unphosphorylated peptide coupled to agarose beads, thus retaining unphosphorylated peptide-specific antibodies. The flowthrough was then loaded onto a column with the phosphorylated peptide coupled to agarose beads, thus retaining the phosphorylated peptide specific antibodies. After elution, the eluate was assayed by enzyme-linked immunosorbent assay against both peptides to control its immunoreactivity and its specificity against the phosphorylation.

### Western Blotting

Mouse brain tissue or cell culture were lysed in 1X sample buffer (Invitrogen) containing 0.1 M DTT. Lysates were sonicated briefly eight times, for the brains, or twice, for the cultures, and denatured at 70 °C for 10 minutes. The samples were centrifuged at 13,300 rpm for 10 minutes and ran on NuPage 4-12 % Bis-Tris polyacrylamide gels (Invitrogen). Proteins were transferred onto an Immobilon PVDF membrane (Millipore), which was then blocked in 4 % milk in tris-buffered saline containing 0.1 % Tween-20 (TBST) for 30 minutes. Primary antibodies were incubated at 4 °C overnight, and HRP-conjugated secondary antibodies at room temperature (RT) for 2 hours. The following primary antibodies were used: rabbit anti-CDKL5 (1:1,000, Atlas HPA002847), mouse anti-tubulin (1:100,000; Sigma T9026), mouse anti-nELAVLs (1:5,000) – Santa Cruz sc-48421) and rabbit anti-p-nELAVLs (1:500). The following secondary antibodies were used at a concentration of 1:10,000: HRP-conjugated anti-rabbit (Jackson 711-035-152) and HRP-conjugated anti-mouse (Jackson 715-035-151). The membrane was developed using ECL reagent (Cytiva) and was visualized with an Amersham Imager 600 or ImageQuant 800 (GE Healthcare). Quantification of Western blots was manually performed using Image Studio Lite Software (version 5.2). nELAVLs phosphorylation was measured relative to total nELAVLs and the other proteins were normalized to tubulin, if not indicated otherwise.

For Phos-tag Western blot, RRM1/2 ELAVl4 samples were lysed in 4X sample buffer containing 0.4 M DTT and heated at 95 °C for 10 minutes. The lysates were then loaded on a Phos-tag SuperSep 12.5% acrylamide 13-well pre-cast polyacrylamide gel (Alphalaboratories) and ran at 20 mA in running buffer containing 25 mM Tris, 192 mM glycine pH 8.3 and 0.1% sodium dodecyl sulfate. The gel was washed for 10 minutes with transfer buffer containing 25 mM Tris, 192 mM glycine pH 8.3 and 10% methanol. Proteins were transferred onto a Immobilon PVDF membrane (Millipore) for 2 hours at 300 mA using transfer buffer. The membrane was then blocked in 4% milk in TBST for 30 minutes, incubated with mouse anti-ELAVL4 (1:5,000 – Santa Cruz sc-48421) at 4 °C overnight and with HRP-conjugated anti-mouse antibody at RT for 2 hours.

### Immunostaining

*Brain slices.* Antigen retrieval was carried out by incubating the brain sections in sodium citrate buffer (10mM tri-Sodium Citrate dihydrate, 0.05% Tween20, pH 6.0) for 20 minutes in the water bath at 95 °C. The sections were let to cool down at room temperature, rinse twice with 0.05% Tween20 in PBS. Then, brain coronal sections were blocked in 10% goat serum with 0.2% Triton x-100 in PBS for one hour at room temperature. Then, the primary antibodies were diluted in the same blocking buffer and incubated overnight at 4 °C in a humid chamber. On the following day, the sections were rinsed three times with PBS prior to the incubation in secondary antibodies diluted in the blocking buffer and counter-stained with DAPI for an hour at room temperature. Afterwards, the sections were rinsed further in PBS and finally mounted in Fluoromount-G (SouthernBiotech, cat. #0100-01) on glass slides. The following antibodies were used: rabbit polyclonal anti-ELAVL2 (Invitrogen, PA5-96475, 1:500), rabbit monoclonal anti-ELAVL3/4 (Abcam, ab184267, 1:500); rat anti-CTIP2 (Abcam, ab18465, 1:1000).

For the imaging of the fluorescent signal, a Leica MPSP8 Upright Confocal microscope equipped with a 63x/1.4 oil or 40x/1.25-0.75 objective and LAS X software used. Images were analysed using Leica LAS X software.

*NSC34 cells.* Cells were transfected 24h prior to seeding on coverslips. After 48h post transfection NSC34 cells were then washed with PBS and fixed using 4% formaldehyde solution (PFA). After permeabilization with 0,5% TRITON X-100, samples stained with DAPI and mounted onto glass slides. Images (GFP and DAPI) were acquired on a Leica-LSM SP8 confocal microscope using a 63x oil objective and the Leica imaging software.

### Rat primary cortical cultures

Cortical neurons from E18 rat embryos were dissected out and plated on 12 mm coverslips at a density of 65,000 cells per well and infected at DIV2 with 4.5 μl of either scrambled-GFP-shRNA or RNY3-GFP-shRNA (AAV2/PHP.eB titer: 1.23 × 10^13^). At DIV9, neurons were treated with either DMSO or 100 nM CAF-382 for 4 hours in a humidified chamber with 5% CO2. Following treatment, cells were incubated with 3 μM puromycin for 10 minutes. As a negative control, 40 μM anisomycin was added 30 minutes prior to puromycin treatment. Incubation was stopped by two washes with prewarmed Neurobasal medium and fixed with 4% paraformaldehyde/sucrose for 12 minutes, followed by immunofluorescence processing. Following permeabilization with 0.25% Triton X-100 and blocking in 1% BSA in PBS, neurons were incubated overnight with rabbit-anti-puromycin (1:3500, Kerafast) rinsed three times with PBS and incubated with the secondary antibody Alexa Fluor 568 rabbit and counterstained with DAPI 1:5000.

AAV vector expressing shRNA under U6 promoter and co-expressing GFP under CMV promoter was designed in VectorBuilder using pscAAV[shRNA]-EGFP-U6 vector backbone. RNY3 shRNA targeting sequence was “AACTAATTGATCACAACCAGT” as shown previously ^67^ and scrambled shRNA sequence was “CCTAAGGTTAAGTCGCCCTCG”, selected from by VectorBuilder recommendations based on no off-targets.

### Mass Spectrometry

#### TMT-labeled quantitative proteomics

##### Tissue Collection and Lysis

Postnatal day 10 mouse brains were collected by cervical dislocation followed by brain dissection. Cerebellum was removed, whole hemispheres were separated and flash-frozen in liquid nitrogen. Frozen tissue was lysed using 8 M urea-based lysis buffer containing 50 mM HEPES (pH 8.2), 10 mM glycerol-2-phosphate, 50 mM NaF, 5 mM sodium pyrophosphate, 1 mM EDTA, 1 mM EGTA, 1 mM sodium vanadate, 1 mM DTT, 1X cOmplete protease inhibitor cocktail (Roche), 1X phosphatase inhibitor cocktail 3 (Sigma), and 500 nM okadaic acid. Protein concentrations were quantified using Pierce BCA assay. Two hundred micrograms per sample were used for TMT 10plex quantitative proteomics.

##### Protein Reduction, Alkylation, and Digestion

Reduction of disulfide bonds was achieved by incubating proteins with 10 mM DTT at 56°C for one hour. Immediately after, samples were alkylated using 20 mM iodoacetamide (IAA) at room temperature under dark conditions for 30 minutes. The reaction was quenched with an additional 20 mM DTT and diluted to reduce the urea concentration to less than 2 M with 50 mM HEPES (pH 8.5). Proteins were then subjected to two-hour enzymatic digestion at 37°C with LysC (4 µg per sample; Lysyl endopeptidase, 125-05061, FUJIFILM Wako Chemicals) followed by overnight digestion with Trypsin (10 µg per sample; MS grade, 90058, ThermoFisher Scientific).

##### Sample Cleanup and TMT Labeling

Post-digestion, peptides were acidified with trifluoroacetic acid (TFA) to a final concentration of 1% and purified using C18-based columns (Nest Group BioPureSPN MACRO; Proto 300 C18; Part# HMM S18V). Samples were subsequently dried in a refrigerated benchtop vacuum concentrator. Peptides were then labeled with TMT 10plex Isobaric Label Reagent Set (0.8 mg per tag, 90110, ThermoFisher Scientific; LOT WD313468) according to the manufacturer’s instructions. Labeling efficiency and mixing accuracy were validated on an Orbitrap Eclipse Tribrid mass spectrometer with a 60-minute high-energy collision dissociation (HCD) MS2 fragmentation method. Labeling efficiency exceeded 99% and mixing check was within 1.5x of total intensity between samples.

##### Phosphopeptide Enrichment and Fractionation

Labeled peptides were quenched with hydroxylamine to a final concentration of 0.2%, pooled, partially dried by vacuum centrifugation, acidified to pH 2.0 with TFA and subjected to cleanup with C_18_ Sep Pak 1cc Vac, 50 mg bed volume (Waters, WAT054955). A final mixing check was performed via LC-MS/MS analysis utilising a 240-minute HCD MS2 fragmentation method. Next, phosphopeptides were captured via Sequential enrichment of Metal Oxide Affinity Chromatography (SMOAC) using high-select TiO_2_ (Thermo Scientific, A32993) followed by Fe-NTA (Thermo Scientific, A32992) columns per manufacturer’s guidelines. Hundred micrograms from the Fe-NTA flowthrough were kept for proteome analysis. Phosphoenriched eluates and proteome sample were dried by vacuum centrifugation, solubilized in 2% TFA, pooled together (phosphoenriched samples), and subjected to high pH reversed-phase fractionation (Thermo Scientific, 84868) using elution solutions for ‘Thermo Scientific TMT-labeled peptides’ (proteome) or ‘unlabelled, native peptides’ (phosphoproteome) as outlined in the manufacturer’s protocol. Fractions were dried by vacuum centrifugation then reconstituted in 0.1% TFA prior to LC-MS/MS analysis.

##### LC-MS/MS Analysis

Phosphoproteomic analyses were performed on an Orbitrap Eclipse Tribrid mass spectrometer. Global phosphoproteome and proteome were analyzed in data-dependent acquisition mode employing 180-minute HCD MS2 and 180-minute multistage activation (MSA) synchronous precursor selection (SPS) MS3 methods as described in ^83^.

##### Data Processing

Raw LC-MS/MS data files from both MS2 and MS3 were co-processed using MaxQuant v2.1.3.0 and v2.3.1.0 against the UniProt mouse reference proteome (UP000000589, canonical November 2019 or with isoforms May 2023). TMT channel intensities were batch-corrected based on manufacturer-supplied factors. Data were searched with Phospho(STY) variable modification, with a false discovery rate (FDR) of 0.01 and a minimum peptide length of 6 amino acids.

##### Statistical Analysis

Processed data were analyzed using an R script based on the ProteoViz ^84^ and Proteus ^85^ packages. The “peptides.txt”, “proteinGroups.txt” and “Phospho(STY)Sites.txt” tables were filtered for reverse hits, potential contaminants and proteins only identified by site. TMT LOT-corrected reporter intensities were normalized using CONSTANd ^86^, log2-transformed, and statistically analyzed using Linear Models for Microarray Data (limma). Significance criteria are reported in the main text and/or figure legends for each experiment. Global protein analysis is presented in the figures and in **Supplementary Data 1**, while differential peptide analysis of the proteome can be found in **Supplementary Data 5**. Additionally, peptide-level expression-change averaging procedure (PECA)^87^ available through Bioconductor (www.bioconductor.org/packages/release/bioc/html/PECA.html) was performed taking the intensity values of each peptide, mean normalizing, log2 transforming, performing statistical test and aggregating into proteins using the leading razor protein. PECA analysis results (**Supplementary Data 5**) largely agree with the summed protein analysis in the main figures.

#### Label-Free Data-Independent Acquisition Mass Spectrometry

##### Sample preparation

Postnatal day 10 and 15 mouse brain tissue was collected and processed as outlined in the TMT-labeled proteomics approach. Two hundred nanograms of digested protein sample were loaded onto an Evotip Pure (desalting C18 trap column; Evosep) per manufacturer’s guidelines. Three injections/tips (technical replicates) were prepared for each sample. Data were obtained employing timsTOF Pro 2 (Bruker) coupled to Evosep One liquid chromatography (LC) system. The LC separation was achieved using the standard 60SPD 2.3 method on an EV1109 column heated to 40°C during analysis. timsTOF Pro 2 was operated in a data-independent acquisition parallel accumulation-serial fragmentation (dia-PASEF) mode with a scan width set to 100-1700m/z with the IM 1K/0 0.6 - 1.6 ramp and accumulation time locked at 100ms. The following windows were used for the data acquisition:

| #MS Type | Cycle Id | Start IM [1/K0] | End IM [1/K0] | Start Mass [m/z] | End Mass [m/z] | CE [eV] |
| --- | --- | --- | --- | --- | --- | --- |
| MS1 | 0 | - | - | - | - | - |
| PASEF | 1 | 0.8400 | 1.6000 | 605.98 | 624.67 | - |
| PASEF | 1 | 0.6000 | 0.8400 | 262.18 | 394.24 | - |
| PASEF | 2 | 0.8700 | 1.6000 | 624.67 | 644.00 | - |
| PASEF | 2 | 0.6000 | 0.8700 | 394.24 | 424.57 | - |
| PASEF | 3 | 0.8900 | 1.6000 | 644.00 | 664.27 | - |
| PASEF | 3 | 0.6000 | 0.8900 | 424.57 | 446.73 | - |
| PASEF | 4 | 0.9000 | 1.6000 | 664.27 | 685.68 | - |
| PASEF | 4 | 0.6000 | 0.9000 | 446.73 | 466.27 | - |
| PASEF | 5 | 0.9000 | 1.6000 | 685.68 | 708.67 | - |
| PASEF | 5 | 0.6000 | 0.9000 | 466.27 | 485.29 | - |
| PASEF | 6 | 0.9100 | 1.6000 | 708.67 | 734.29 | - |
| PASEF | 6 | 0.6000 | 0.9100 | 485.29 | 503.27 | - |
| PASEF | 7 | 0.9200 | 1.6000 | 734.29 | 763.37 | - |
| PASEF | 7 | 0.6000 | 0.9200 | 503.27 | 520.81 | - |
| PASEF | 8 | 0.9400 | 1.6000 | 763.37 | 797.43 | - |
| PASEF | 8 | 0.6000 | 0.9400 | 520.81 | 537.63 | - |
| PASEF | 9 | 0.9500 | 1.6000 | 797.43 | 838.95 | - |
| PASEF | 9 | 0.6000 | 0.9500 | 537.63 | 554.30 | - |
| PASEF | 10 | 0.9600 | 1.6000 | 838.95 | 892.42 | - |
| PASEF | 10 | 0.6000 | 0.9600 | 554.30 | 571.49 | - |
| PASEF | 11 | 0.9800 | 1.6000 | 892.42 | 971.97 | - |
| PASEF | 11 | 0.6000 | 0.9800 | 571.49 | 588.73 | - |
| PASEF | 12 | 1.0400 | 1.6000 | 971.97 | 1199.58 | - |
| PASEF | 12 | 0.6000 | 1.0400 | 588.73 | 605.98 | - |

##### Data processing

Raw data from timsTOF Pro 2 dia-PASEF method was processed using DIA-NN^88^ (Data-Independent Acqusition by Neural Networks) version 1.8.1 against *in silico* generated spectral library of UniProt mouse reference proteome (UP000000589, March 2021). Default settings were used with a few modifications as follows: data were searched with tryptic digest missed cleavage of 2 and match between runs option on. False-discovery rate was set to 0.01.

##### Data analysis

The “report.tsv” file was imported into DIAgui^89^ (a Shiny application for processing output data from DIA-NN). Peptides and Protein groups files were obtained using the fast MaxLFQ algorithm from iq package. Default settings in DIAgui were used including Filter q-values for peptides and protein groups were set at 0.01 and “Only keep peptides counts all” (proteins) and “Proteotypic only” (peptides and proteins) options were selected. Processed data were further analysed by excluding any peptides/proteins that have missing value(s) for any of the mass spectrometry runs. Technical repeats (injections) were averaged, and data were statistically analysed using the *limma* package. Differential protein analysis can be found in the figures and in **Supplementary Data 4**, while differential peptide analysis can be found in **Supplementary Data 5**.

Additionally, peptide-level expression-change averaging procedure (PECA) available through Bioconductor (www.bioconductor.org/packages/release/bioc/html/PECA.html) was performed taking the peptides table with log2 transformed and MaxLFQ (iq package) normalized intensity values from DIAgui. Peptides that contained missing values were removed and technical replicates were averaged. Normalized, log2 transformed and filtered data was then subjected to PECA statistical analysis and protein aggregation using the gene name. PECA analysis results (**Supplementary Data 5**) largely agree with the protein analysis in the main figures.

### Protein purification

*His-CDKL5*. Purification of His6-CDKL51–352 (kinase domain) was performed using infected Sf9 insect cells using pFastBacHT-His6-CDKL51–352. 1 L of Sf9 insect cells at 2 × 106 cells/mL was infected with 2 mL of P3 for 48 hours and centrifuged at 3,000 g for 15 minutes at 4 °C. Cell pellets were resuspended in 25 mM NaHCO3 and lysed with 2X lysis buffer containing 100 mM Tris–HCl pH 7.5, 1 M NaCl, 30 mM imidazole pH 8.0, 5% glycerol, 10 mM MgSO4, 0.3% Triton X-100, 2 mM TCEP, 1X protease inhibitor cocktail (Roche) and 1:1,000 nuclease. Lysate was centrifuged at 55,000 g for 30 minutes at 4 °C. Supernatant containing solubilised His6-CDKL51–352 was loaded on HisPur cobalt resin (Thermo Scientific) and washed three times with wash buffer containing 50 mM Tris–HCl pH 7.5, 500 mM NaCl, 5% glycerol, 1 mM TCEP, 0.01% Triton X-100 and 15 mM imidazole. Bound proteins were eluted in 1 mL wash buffer containing 500 mM imidazole. Protein concentration was measured by comparison with bovine serum albumin standards.

*mELAVL4 (RRM1/2)*. T7 Express lysY Competent bacteria (NEB) were transformed with pET47b-His6-mELAVL414-215 plasmid using manufacturer’s instructions. The bacteria were then grown at 37 °C on a shaker, induced with 0.4 mM of isopropyl β-D-1-thiogalactopyranoside for another 3 hours. Bacteria pellet was lysed in 30 mL of loading buffer containing 50 mM Tris pH 8.0, 300 mM NaCl, 0.5% Triton X-100, 1X protease inhibitor cocktail (Roche) and 2 mM PMSF. The lysate was sonicated 2 to 6 times at 15 A, rotated at 4 °C for 30 minutes and centrifuge at 4 °C at maximum speed for 10 minutes. Supernatant containing solubilised His6-mELAVL414-215 was loaded on HisPur beads (Thermo Scientific) and washed once with the loading buffer, twice with wash buffer 1 (50 mM Tris pH 8.0, 1 M NaCl, 10 mM imidazole) and wash buffer 2 (50 mM Tris pH 8.0 and 150 mM NaCl). The beads containing His6-mELAVL414-215 were then incubated overnight with 360 μL of HRV 3C protease (Thermo Scientific) in 9 mL of wash buffer 2 in order to cleave the His6 tag from the protein. The beads were centrifuged at 500 g for 4 minutes at 4 °C and the supernatant was loaded on fresh HisPur beads (Thermo Scientific) as an additional cleaning step. The beads were again centrifuged and the supernatant containing purified untagged mELAVL414-215 was transferred to a new tube. The purified protein was finally concentrated using a Vavaspin 20, 3,000 MWCO PES column (Sartorius) and its final concentration was measured using Bradford assay (Thermo Scientific).

### *In vitro* kinase assay

Purified His-CDKL5 and RRM1/2 ELAVL4 were incubated with kinase buffer containing 20 mM Tris-HCl pH 7.5, 10 mM MgCl2, 1X protease inhibitor cocktail (Roche), 1 μM okadaic acid (Fluorchem), 1 mM DTT and 0.5 mM ATP at 30 °C for 4 hours. The reaction was then incubated for 1 hour on HisPur beads (Thermo Scientific) and centrifuged at 500 g for 3 minutes at 4°C to remove CDKL5 from the preparation. The supernatant containing purified phosphorylated RRM1/2 ELAVL4 was finally dialysed and transfer to a final buffer with 50 mM Tris pH 8.0, 150 mM NaCl and 0.5 mM TCEP. The phosphorylation level of the final purified was assessed using Phos-tag gel and intact mass spectrometry. The same preparation was used for the bio-layer interferometry (BLI) assay.

### Bio-layer interferometry (BLI)

BLI assay was performed by Laura Masino from the Structural Biology facility at the Francis Crick Institute, using the Octet RED96 as described previously ^90^. In short, using a sensor rack and 96-well microplates, the 8-channel streptavidin-coated biosensors were soaked (1) in experimental buffer (baseline 1), (2) in experimental buffer containing 0.5 μg/μL of RNA probes (RNA probe binding), (3) in experimental buffer again (baseline 2), (4) in experimental buffer containing purified RRM1/2 ELAVL4 in different concentrations from 1 μM to 8 nM with one concentration per channel (protein binding) and (5) in experimental buffer (dissociation). Binding dynamics of unphosphorylated RRM1/2 ELAVL4 to a control probe (biotin-GC(x10)) and an experimental probe (biotin-U(x20)) were first compared. Then binding dynamics between unphosphorylated and phosphorylated RRM1/2 ELAVL4 to the experimental probe were compared. The data was analysed using an in-house software and the association amplitude binding curve was fitted using non-linear least squares methods.

### iCLIP

Fresh brains were taken promptly from 5 *Cdkl5* WT and 5 *Cdkl5* constitutive KO mice at P10 after cervical dislocation and decapitation. The cerebellum was dissected out of the brains and the hemispheres were separated. The two hemispheres were then flash frozen separately in liquid N_2_. One hemisphere of each animal was powdered in liquid N_2_ using a pestle and mortar. Half of the powder was dedicated to improved individual nucleotide resolution crosslinking and immunoprecipitation (iCLIP) and the other half to 3’ mRNA sequencing.

Improved iCLIP (iiCLIP) was performed as described previously ^61^. In short, frozen brain powder were ultraviolet crosslinked 4x with 150 mJ/cm2 on dry ice using a Stratalinker 2400 (Stratagene) at 254nm and lysed with 1ml lysis buffer (50 mM Tris-HCl, pH 7.4, 100 mM NaCl, 1% Igepal Ca-630 (Sigma I8896), 0.1% SDS, 0.5% sodium deoxycholate) supplemented with 1X protease inhibitor cocktail (Roche), 1:100 phosphatase inhibitor III (Roche), 1X Pierce phosphatase Inhibitor (Thermo Scientific), and 0.5 μM of okadaic acid (Fluorchem). Samples were then transferred to 1.5 ml Bioruptor® microtubes (Diagenode, C30010016) and sonicated for 10 cycles with alternating 30 sec on/off at low intensity using Bioruptor. Lysate concentration was determined using Biorad DC assay and diluted to 1 mg/ml. One millilitre of each sample was then partially digested with 0.4 units/ml RNase I (Thermo Scientific, EN0602) and 2 μl Turbo DNase (Ambion, AM2238) for 3 min at 37°C shaking at 1100 rpm followed by immediate transfer to ice. Cell lysates were cleared by centrifugation for 10 min at 4°C at 21,000 xg followed by Proteus Clarification Mini Spin Column. nELAVLs–RNA complexes were immunoprecipitated with an antibody against ELAVLs (Santa Cruz – sc-5261) pre-immobilised on protein G Dynabeads (Dynal) rotating end-over-end overnight at 4°C. Additionally, two IgG controls were added where normal mouse IgG antibody (Santa Cruz, sc-2025) was used, instead of anti-ELAVLs antibody, for 1 WT and 1 KO. Beads were washed twice with high-salt wash buffer (50 mM Tris-HCl, pH 7.4, 1M NaCl, 1 mM EDTA, 1% Igepal CA-630, 0.1% SDS, 0.5% sodium deoxycholate) and once with PNK buffer (20 mM Tris-HCl, pH 7.4, 10 mM MgCl_2_, 0.2% Tween-20) before 3’ end RNA dephosphorylation. Beads were then resuspended in buffer containing 8 μl 5x PNK pH 6.5 buffer (350 mM Tris-HCl, pH 6.5, 50 mM MgCl_2_, 5 mM DTT), 1 μl PNK (NEB M0201L), 0.5 μl FastAP alkaline phosphatase (ThermoFisher, EF0654), 0.5 μl RNasin and 30 μl H_2_O, and incubated for 40 min at 37°C shaking at 1100rpm. Following a wash in ligation buffer (50 mM Tris-HCl, pH 7.4, 10 mM MgCl_2_, 1 mM DTT), RNA was ligated at their 3′ ends to an RNA L3 infrared adaptor to allow visualisation. Beads were resuspended in 6.3 μl H_2_O, 3 μl 10x DTT-free ligation buffer (500 mM Tris-HCl, pH 7.5, 100 mM MgCl_2_), 0.8 μl 100% DMSO, 2.5 μl T4 RNA ligase I (NEB, M0437M), 1.4 μl RNasin, 1.5 μl PNK (NEB, M0201L), 2.5 μl pre-adenylated adaptor L3 (stock 1 μM), 9 μl 50% PEG8000 and incubated overnight at 16°C shaking at 1100 rpm. Adaptor ligated samples were washed twice with high-salt wash buffer followed by a PNK buffer wash. The beads were then resuspended in the following reaction for adapter removal: 12.5 μl H_2_O, 2 μl NEB Buffer 2, 0.5 μl 5’ Deadenylase (NEB, M0331S), 0.5 μl RecJ_f_ endonuclease (NEB, M0264S), 0.5 μl RNasin, 4 μl 50% PEG8000 and incubated for 1 hour at 30°C followed by incubation at 37°C for 30 min shaking at 1100 rpm. Following two washes in high-salt wash buffer and two washes in PNK buffer, samples were eluted using 1x NuPAGE LDS loading buffer supplemented with DTT and used for gel electrophoresis and nELAVLs-RNA-L3 transfer onto a Protan BA85 nitrocellulose membrane. The membrane was then visualised using a LI-COR Odyssey-Clx. Crosslinked nELAVLs-RNA-L3 complexes were excised from the membrane and protein was removed with 10 μl proteinase K (Roche, 03115828001) and 190 μl PK_SDS buffer (10 mM Tris-HCl, H 7.4, 100 mM NaCl, 1mM EDTA, 0.2% SDS) shaking for 60 min at 50°C followed by phenol/chloroform RNA isolation. RNA was then precipitated overnight at -20°C in the presence of 0.5 pmol barcoded primer, glycoblue (Ambion), NaAc and EtOH. Samples were then centrifuged at 21,000 xg for 30 min, pellets were washed once with 80% EtOH and centrifuged again before pellet resuspension in 5.5 μl H_2_O. The purified RNA was then reverse transcribed with 0.5 pmol of the same barcoded reverse transcription primer using the Superscript IV Kit (Thermo Scientific) and following manufacturer’s instructions. Template RNA was removed by alkaline hydrolysis using NaOH for 15 min at 85°C followed by neutralisation with HCl. cDNA molecules were purified using AMPure XP beads (Beckman Coulter) following manufacturer’s protocol and eluted in 6.75 μl H_2_O, then circularised in buffer containing 3 μl 5x TS2126 ligase buffer, 0.75 μl 1 mM ATP, 0.75 μl 50 mM MnCl_2_, 3 μl 5 M betaine, 0.75 μl TS2126 RNA ligase (purified in-house) and incubated at 60°C overnight. Circularised cDNA was purified again with AMPure XP beads (Beckman Coulter) and eluted 10 μl. cDNA was then PCR-amplified using P5Solexa/P3Solexa primer mix (Solexa, 10 μM each) and Phusion HF Master mix (NEB) for 16-18 cycles and purified again using AMPure XP beads. Purified cDNA libraries were size-selected (145-400 nt) by gel purification (6% TBE), excision, extraction and overnight precipitation in the presence of glycoblue, NaAc and EtOH. High-quality cDNA libraries were pooled together and subjected to high-throughput 100bp single-read sequencing using NovaSeq (Illumina). The sequencing was carried out by the Advanced Sequencing Facility at the Francis Crick Institute.

#### Analysis

Raw reads were processed using the clipseq nfcore pipeline (version 1.0.0). The pipeline uses the nf-core framework^91^ with nextflow (version 21.04.0)^92^. In short, reads were trimmed using cutadapt (version 3.0)^93^ and subsequently pre-mapped to rRNA and tRNA with Bowtie2 (version 2.4.2)^94^. The remaining unmapped reads were aligned against the mouse genome GRCm38 and annotation release 95, both from Ensembl, with STAR (version 2.6.1d)^95^. Peak calling was performed with Paraclu (version 9)^96^ with a minimum score of 10, a maximum cluster length of 200 and a minimum density increase of 2.

Consensus peak annotation and initial quality assessment was performed in R-4.0.0 (R Core Team, 2020) using the Bioconductor packages RCAS (version 1.16.0)^97^ and plyranges (version 1.10.0)^98^.

Consensus peaks were defined as those peaks that are present in at least four samples across the ten non-IgG samples. For peak-to-gene annotation we only considered peaks falling into those annotated as type == “gene” and removed those that fall on the mitochondrial genome. Additionally, we removed peaks from the subsequent analysis that overlap more than one annotated gene. The overlap between consensus peaks and genes was considered sufficient if the peak only partially overlapped the annotated gene. Consensus peaks were annotated to transcript features with the following precedence: threeUTRs > fiveUTRs > promoters > exons > introns > cds > transcripts.

DREME^99^ is used for basic motif calling using peaks. As input for DREME the sequence from the region +/-20nt around the crosslink site is used as input for DREME.

Differential binding analysis was performed in R-4.0.0 (R Core Team, 2020) using the DESeq2^100^ package (version 1.30.1). The input count matrix contains the number of crosslink sites within each consensus peak for each sample. Using the model ∼genotype, differential ELAVL-binding was compared between CDKL5 knock-out and control genotypes at postnatal day 10 (P10). Differentially bound consensus peaks were identified as those with an FDR < 0.05 (Benjamini & Hochberg adjustment) and their regularised (“shrunken”) log2 fold changes were provided.

For peak visualization, .bam files from the same genotype were merged using samtools. Peaks were visualized using *clipplotr*^101^.

### RNA isolation and sequencing library preparation

Total RNA from brain tissue was isolated using the RNeasy Plus Mini kit from Qiagen following manufacturer’s instructions.

#### Library preparation for bulk RNA-seq

Sequencing libraries were prepared using the KAPA mRNA HyperPrep Kit (Roche, kit code KK8581) according to manufacturer’s instructions. Briefly, samples were normalised to 1000 ng of RNA in a final volume of 50 µl. PolyA-tailed RNA was captured with 50 µl of capture beads at 65 °C for 2 min and 20 °C for 5 min. Beads were washed to remove other RNA species and RNA was eluted from the beads in 50 µl of RNase free water by incubating at 70 °C for 2 min and 20 °C for 5 min. A second capture of PolyA-tailed RNA was performed by adding 50 µl of bead binding buffer and the sample incubated for 5 min at 20 °C.

Beads were washed to remove other RNA species and eluted in 20 µl of Fragment, Prime and Elite Buffer. The fragmentation reaction was run for 6 min at 94 °C for a library insert size of 200-300 bp. Fragmented RNA underwent first and second strand cDNA synthesis according to manufacturer’s instructions.

The KAPA Unique Dual-Indexed Adapters Kit (15μM) (Roche- KK8727) stock was diluted to 7 µM as recommended by the manufacturer for a total RNA input of 1000 ng. The adaptor ligation was carried out by adding 5 µl of diluted adaptor and 45 µl of ligation mix (50 µl of ligation buffer + 10 µl of DNA ligase) to 60 ul of cDNA. The ligation reaction was run for 15 mins at 20°C.

To remove short fragments such as adapter dimers, two SPRISelect bead clean-ups were done (0.63x SPRI and 0.7x SPRI). To amplify the library, 25 µl of Kapa HiFi HotStart PCR master mix plus 5ul of Library Amplification Primer mix was added to 20 µl cDNA and 8 PCR cycles were performed as recommended by the manufacturer for a total RNA input of 1000 ng. Amplified libraries were purified via a SPRISelect 1x bead cleanup.

The quality and fragment size distributions of the purified libraries was assessed by a 4200 TapeStation Instrument (Agilent Technologies).

Libraries were pooled and sequenced on the Illumina NovaSeq6000 in PE100 configuration to an average read depth of 25M.

#### Library preparation for 3’Sequencing

3’ RNA-seq libraries were prepared using Quantseq FWD kit (Lexogen) and 100 ng RNA as input following manufacturer’s protocol. The quality of the purified libraries was assessed using an Agilent D1000 ScreenTape Kit on an Agilent 4200 TapeStation. Libraries were sequenced to a depth of at least 25M reads on an Illumina NovaSeq 6000 run in 101-8-8-101 configuration.

### Bulk RNA sequencing and analysis

#### *Cdkl5* KO mice

Raw reads were processed using the RNA-seq nfcore pipeline (version 3.5): Patel, H 2021. nf-core/rnaseq: nf-core/rnaseq v3.2 - Copper Flamingo. (2021) doi:10.5281/zenodo.4984258. The pipeline uses the nf-core framework ^102^ with nextflow (version 21.10.6) ^103^. In short, reads were trimmed using trimgalore (version 0.6.7) (Krueger et al. FelixKrueger/TrimGalore: v0.6.10 - add default decompression path (0.6.10). Zenodo. https://doi.org/10.5281/zenodo.5127898) and subsequently aligned and quantified with STAR-RSEM (version 1.3.1) ^95, 104^ against the mouse genome GRCm38 and annotation release 95, both from Ensembl.

Differential gene expression analysis was performed in R-4.0.0 (R Core Team, 2020) using the DESeq2 ^105^ package (version 1.30.1). Using the model ∼genotype the gene expression between CDKL5 knock-out and control genotypes was compared at postnatal day 10 (P10). The regularised (“shrunken”) log2 fold changes (lfc) for genes of interest were visualised as a heatmap using the ComplexHeatmap ^106^ package (version 2.6.2). Significance criteria are reported in the main text and/or figure legends.

Hom *Elavl2/3* SA mice raw reads were processed in the web-based platform https://app.flow.bio/ using the RNA-seq pipeline version 3.12 based on nf-core/rnaseq v3.12.0 and nextflow version 23.04.3. Briefly, reads were trimmed using trimgalore, aligned and quantified using star_rsem against the mouse reference proteome GRCm38/mm10 release 102.

Differential gene expression analysis was performed in RStudio using R and DESeq2 statistical analysis. Animals were compared using the model ∼genotype. Shrunken log2 fold change was calculated. Significance criteria are reported in the main text and/or figure legends.

### Alternative splicing (AS) from bulk RNA-sequencing

<u>Whippet (</u>https://github.com/timbitz/Whippet.jl) was used for the identification of alternative splice events (whippet-quant.jl). To identify splice events which are differentially regulated between conditions whippet-delta.jl was used. The analysis of differentially spliced nodes was performed with the following parameters: 1) Whippet was run with bias correction – 1.1) accounts for 5’sequence bias using the method of ^107^ in which all read counts are adjusted after read-alignment but before any quantification takes place; 1.2) for alternative splicing quantification only, Whippet will additionally correct for GC-content bias using a fast heuristic method; 2) Whippet-delta settings used were – 2.1) minimum samples: 4; 2.2) minimum reads: 10; 3) for individual samples: 0.1 < Psi < 0.9; 4) for the identification of differentially alternatively spliced nodes – 4.1) probability >= 0.9; 4.2) |deltaPsi| > 0.1 (at least 10% change); 4.5) confidence interval (CI) < 0.2 to ensure that events supported by low numbers of mapped reads (and therefore high variability) are excluded from further analysis.

### PolyA site usage analysis (APA) from 3’sequencing

Reads were pre-processed and mapped against the GRCm38 Mus musculus genome (with the GENCODE version M20 annotation) using the nf-quantseq pipeline (v.0.1.0; DOI: 10.5281/zenodo.14417140). This pipeline also generates a pA site atlas as described in Hallegger et al. ^108^, as well as tallies the number of reads mapping to a window 200 nucleotides upstream of each poly(A) site in a strand-aware manner. This count table served as input for DRIMSeq ^109^. Only genes with at least 10 reads in 75% of all samples, and PAS with at least 5 reads in 75% of samples in the smallest group size were retained for the DRIMseq analysis. DRIMseq was used to fit gene-level Dirichlet-multinomial models and feature-level beta-binomial models. A likelihood ratio test assessed differential PAS usage between conditions. We employed a two-stage testing procedure using the stageR package^110^. to control for multiple testing and maintain a gene-level false discovery rate of 5%. PAS with adjusted p-values less than 0.05 and that exhibited an absolute change in usage of at least 10% between conditions were deemed significantly altered pA sites. The code used to run the DRIMseq package can be viewed on Github (https://github.com/ulelab/DRIMseq_script_Ultanir) and is also archived on Zenodo (https://doi.org/10.5281/zenodo.15043754).

### Protein synthesis measurement by immunostaining

To measure the protein synthesis in primary cortical neurons (DIV9) transfected with GFP we followed an established protocol ^69^. Briefly, neurons were treated with DMSO or 100nM CAF-382 for 4 hours in conditioning medium and kept in a humidified incubator at 37C with 5% CO2 to inhibit CDLK5 activity. Neurons were then incubated with 3 µM Puromycin for 10 minutes to label the newly synthesized proteins. To validate the detected signal of puromycin-labelled protein by anti-puromycin antibody, negative control experiment neurons were pre-treated with 40 µM anisomycin in conditioning medium for 30 minutes prior to puromycin treatment to inhibit puromycin incorporation. After the incubation, neurons were washed twice with pre-warm PBS before fixation with 4% PFA/Sucrose for 12 minutes. After fixation samples were processed for immunofluorescence and puromycin signal was quantified in the soma and in 30 µm of dendritic length.

### Neuropixels recordings

#### Surgical procedures

Mice underwent two surgeries (metal headplate implantation and craniotomy). Analgesia was provided on the day prior to surgery (Metacam 5mg/kg in custard). For both surgeries, mice were anesthetised with isoflurane (5% induction, 2-2.5% maintenance) and injected subcutaneously with analgesic and anti-inflammatory compound (10 mg/kg Meloxicam and 0.1 mg/kg Buprenorphine, dosage of injectable 0.1 ml/10g of body weight). The animals body temperature was maintained at 37 C° with a thermal mat. Animals were provided analgesics (Metacam 5mg/kg in custard) for 48hs after surgery. A metal headplate was fixed to the skull using dental cement (Super Bond C&B, Sun Medical) centred on the right primary visual cortex (V1) (AP: 0.4 mm and ML: 2.7 mm from bregma). After one week of habituation to the behavioural apparatus animals underwent a second surgery to perform a craniotomy of 2-3 mm in diameter and a durectomy. The craniotomy was covered with artificial dura (Dura-Gel, Cambridge NeuroTech) and sealed with Kwik-Cast silicone elastomer (World Precision Instruments). Mice were left to recover for at least 24 hours before recordings.

#### Recordings

After recovery, neural activity was recorded using Neuropixels 1.0 ^111^. Neuropixels recordings were performed on awake, head-restrained animal using a National Instruments I/O PXIe-6341 module and OpenEphys software (https://github.com/open-ephys). The probe was coated with DiI (ThermoFisher Scientific, V22885) or DiO (ThermoFisher Scientific, V22886) on the day of the recording, to be able to validate recording location. On the last day of recordings, if brain tissue was not required for other experiments, the animals were perfused and the extracted brains were imaged with serial two-photon (STP) tomography ^112^ to perform post-hoc tracing of the electrode locations. Brainreg was used to register the brains to a reference atlas (atlas.brain-map.org) and the probes were traces with brainreg-segment^113^. During the recording, the mouse was allowed to run freely on a cylindrical treadmill made of polystyrene foam.

Sessions were automatically spike-sorted using Kilosort2 (https://github.com/MouseLand/Kilosort/releases/tag/v2.0) and manually curated to select isolated single cells using Phy (https://github.com/cortex-lab/phy).

#### Stimuli

Visual stimuli were generated in Matlab using Psychophysics Toolbox ^114^ and presented on a calibrated LCD monitor (60 Hz refresh rate) positioned 20 cm from the left eye at approximately 45° to the long axis of the animal, covering ∼110° × 80°of visual space. Stimuli for orientation selectivity mapping consisted of square wave gratings (0.03 cycles per degree, measured at the shortest distance between the eye and the monitor, 2 Hz, 100% contrast) drifting in 16 different directions for 1.5 s were presented randomly and were interleaved with a grey screen (∼1 s) between grating presentations. Each grating direction was repeated 12 times.

Receptive field mapping stimuli consisted of black (<0.05 cd m-2) and white (43 cd m-2) squares of 9.6° × 7° on a grey background (23 cd m-2). The squares were presented one at a time and in random order at one of 80 positions (10 × 8 matrix covering a total area of 96° × 70°; each position was repeated 12 times). The presentation rate was ∼1.7 Hz and the duration of each stimulus was ∼0.15 s, followed by 0.75 s blank screen.

#### Data analysis

All analyses were performed in Matlab (MathWorks). Spontaneous firing was determined during blank stimuli of 5 minutes duration in between visual stimulation protocols.

Power spectrum analysis on the local field potential (LFP) activity was performed using the Welch’s method and power values were calculated for specific frequency bands, including delta (1-4 Hz), theta (4-8 Hz), alpha (8-12 Hz), beta (12-30 Hz), and gamma (30-100 Hz).

To obtain a smooth orientation tuning curve, Fourier interpolation was applied to the averaged response to different orientations. The preferred orientation was determined as the minimum angle between the peak and the null direction. The response magnitudes at these preferred and null orientations are extracted from the interpolated tuning curve.

Orientation selectivity was estimated with the orientation selectivity index (OSI). The formula for calculating OSI is:

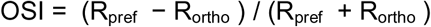

Where R_pref_ is the response magnitude at the preferred orientation, while R_ortho_ is the response to the orthogonal orientation. The OSI ranges from 0 to 1, where higher values indicate a higher orientation selectivity.

Direction selectivity index (DSI) was calculated as below:

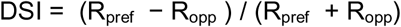

Where R_pref_ is the response magnitude at the preferred orientation and direction, while R_opp_ is the response to the opposite direction. The DSI ranges from 0 to 1, where higher values indicate a higher direction selectivity.

The ON and OFF spatial RFs were obtained separately by analysing the responses to white and black stimulus patches. The Wilcoxon signed-rank test was used to assess responsiveness to individual locations, comparing baseline FR (400 ms prior to the stimulus) with stimulus-evoked FR (400 ms after the stimulus). Benjamini-Hochberg procedure was used to perform multiple comparison correction. Neurons that did not pass this test were excluded from further analysis. We obtained combined ON and OFF RF subfield maps of visually tuned neurons by fitting a two-dimensional Gabor function using the Levenberg–Marquardt algorithm.

For onset responses, the interval of interest spanned from 50 ms to 250 ms post-stimulus presentation. The mean response profile for each neuron was quantified through the calculation of the area under the curve (AUC). Mean firing rate during offset responses were calculated for the time interval between 250 and 400 ms following stimulus presentation. To normalise responses the firing rate of individual neurons was divided by their baseline activity immediately preceding the stimulus presentation (400 msec).

To map the ON and OFF subfields of individual neurons we averaged across repetitions for each location the firing rate for the first 400 ms of the stimulus presentation. We fitted the resulting 2D map using an elliptic 2D Gaussian and then determined the RF area from the fit. Only neurons with an R-squared greater than 0.5 were included in this analysis.

The RF selectivity index was calculated as:

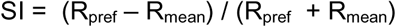

Where R_pref_ is the average response magnitude at the preferred retinotopic location across 12 repetitions, while R_mean_ is the mean response across all locations tested. The SI ranges from 0 to 1, where higher values indicate a higher degree of selectivity.

### Statistics

Data were analysed using Graphpad Prism 10 (Graphpad software, Inc., San Diego, CA, USA). Statistical significance between groups was determined by one- or two-way ANOVA followed by pairwise comparisons or two-tailed unpaired Student t test as indicated in the figure legends. Statistical significance was visualised as: ns=not significant (p>0.05), *=p<0.05, **=p<0.01 and ***=p<0.001.

For Neuropixels data analysis to test for statistical significance between the two groups we used non-paired non-parametric tests (Mann–Whitney U test).

## Data availability

The mass spectrometry proteomics data have been deposited to the ProteomeXchange Consortium (http://proteomecentral.proteomexchange.org) via the PRIDE partner repository^115^ with the dataset identifiers PXD058739, PXD058776. The RNA sequencing data discussed in this publication have been deposited in NCBI’s Gene Expression Omnibus and are accessible through GEO Series accession numbers GSE289273, GSE289015, GSE288923, GSE288922, GSE294025. Western blot images, immunostaining images and spike sorted data and LFP traces will be made publicly available at Figshare.

## Code availability

R code for proteomics work will be made available on Github/Zenodo upon publication. Python code for analysing the data and generating Fig. 6 will be made available on Github upon publication.

## Acknowledgements

We thank the Francis Crick Institute Genetic Modification Services (GeMS) for generating *Elavl2/3/4* phosphomutant mice. We thank Laura Masino at the Francis Crick Institute Structural Biology technology platform for BLI experiments; Prof. Alysson Muotri (UCSD) for human iPSC derived neuron lysates. This work was supported by the Francis Crick Institute which receives its core funding from Cancer Research UK (CC2037), the UK Medical Research Council (CC2037), and the Wellcome Trust (CC2037) and project grants from the Loulou Foundation. For the purpose of Open Access, the author has applied a CC BY public copyright license to any Author Accepted Manuscript version arising from this submission.

## Ethics Declarations

Authors declare no conflict of interest.

