## Supplementary Figure 1 for "CDKL5 phosphorylates neuronal ELAVL proteins to promote mRNA binding, protein synthesis and visual cortex development"

**a**

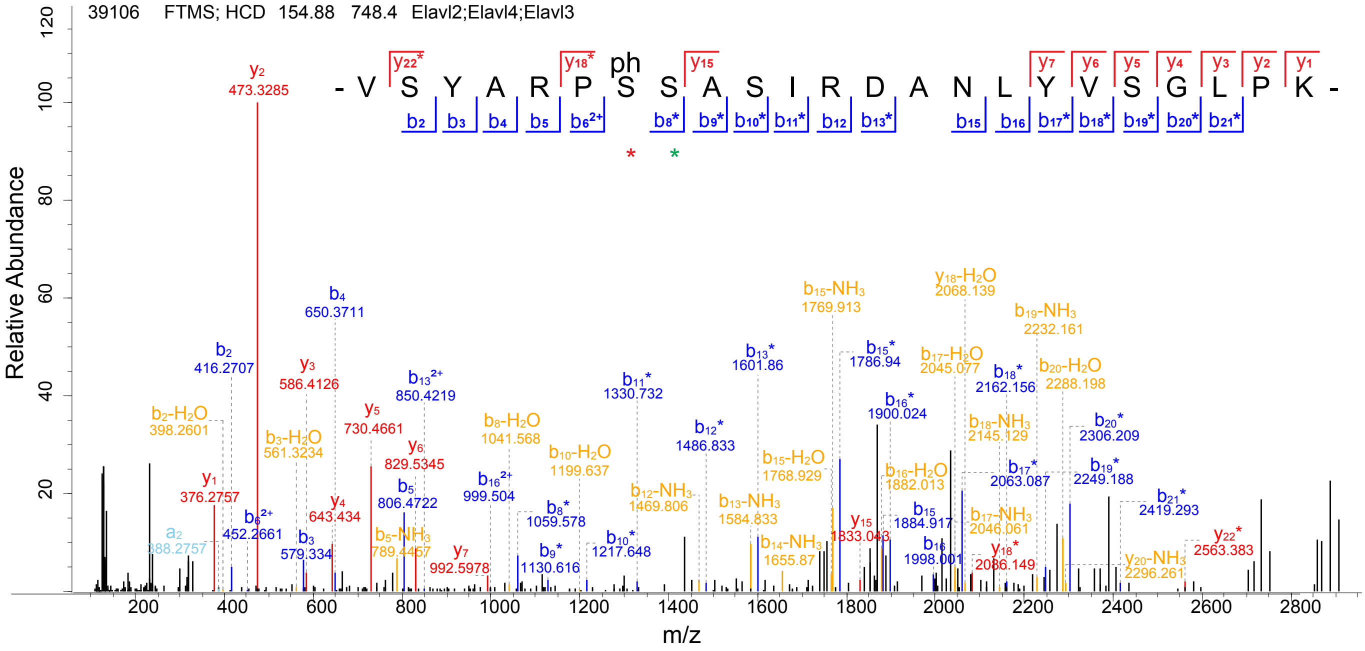


**Supplementary Fig. 1: Annotated MS2 fragmentation spectrum of the TMT labelleled phosphopeptide "VSYARPSSA-SIRDANLYVSGLPK".**

**a** Best localized spectrum for phosphopeptides in ELAVL2,3, and 4 (corresponding to UniProt IDs: Q60899, Q60900 and Q61701, respectively) acquired in a TMT-labelled mass spectrometry approach when comparing *Cdkl5* full WT and KO mouse brains. Spectrum was exported from the Viewer section of MaxQuant software.
