## Supplementary Figure 2 for "CDKL5 phosphorylates neuronal ELAVL proteins to promote mRNA binding, protein synthesis and visual cortex development"

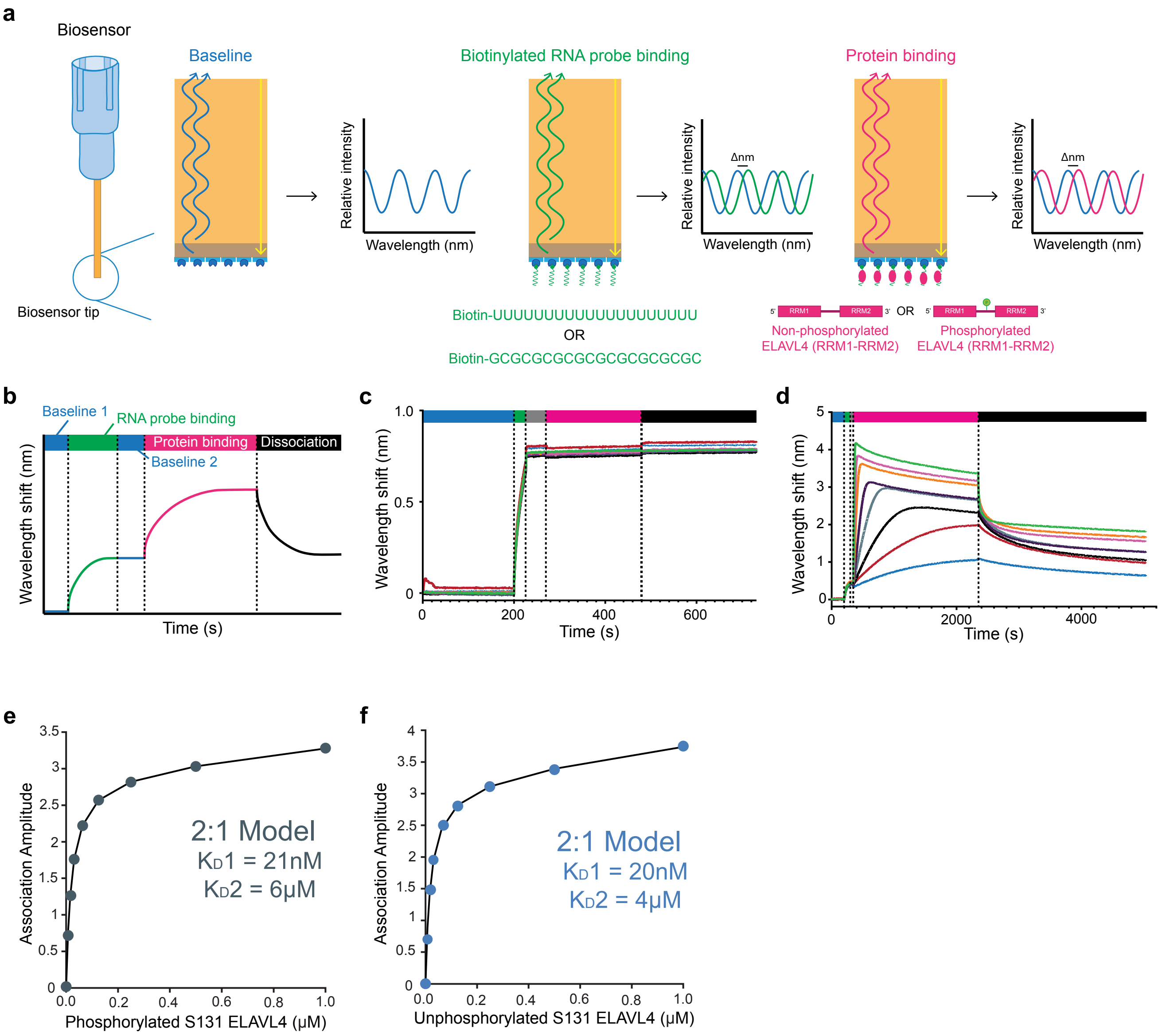


**Supplementary Fig. 2: ELAVL4 binding dynamics to U-rich RNA is not changed in absence of Ser 131 phosphorylation *in vitro.***

**a** Schematic of bio-layer interferometry quantifying binding kinetics between RNA probes and RNA-binding protein by measuring the shifts in reflected white light upon changes in thickness of the streptavidin biolayer. **b** Schematic of a wavelength shift generated by a typical BLI experiment with the wash of the biosensor tip with the main buffer (baseline 1), the binding of the RNA probe to the tip (RNA probe binding), the second wash with the main buffer (baseline 2), the binding of the protein to the RNA probe (protein binding) and the dissociation between the protein and the RNA probe (dissociation). **c** The wavelength shift generated by the binding of different concentrations of ELAVL4 (RRM1-RRM2) to the control probe biotin-GC(x10). **d** The wavelength shift generated by the binding of different concentrations of ELAVL4 (RRM1-RRM2) to the experimental probe biotin-U(x20). **e** Association amplitude binding curve between phosphorylated Ser 131 ELAVL4 and biotin-U(X20) probe and its best fitted model. **f** Association amplitude binding curve between unphosphorylated Ser 131 ELAVL4 and biotin-U(X20) probe and its best fitted model.
