## Supplementary Figure 3 for "CDKL5 phosphorylates neuronal ELAVL proteins to promote mRNA binding, protein synthesis and visual cortex development"

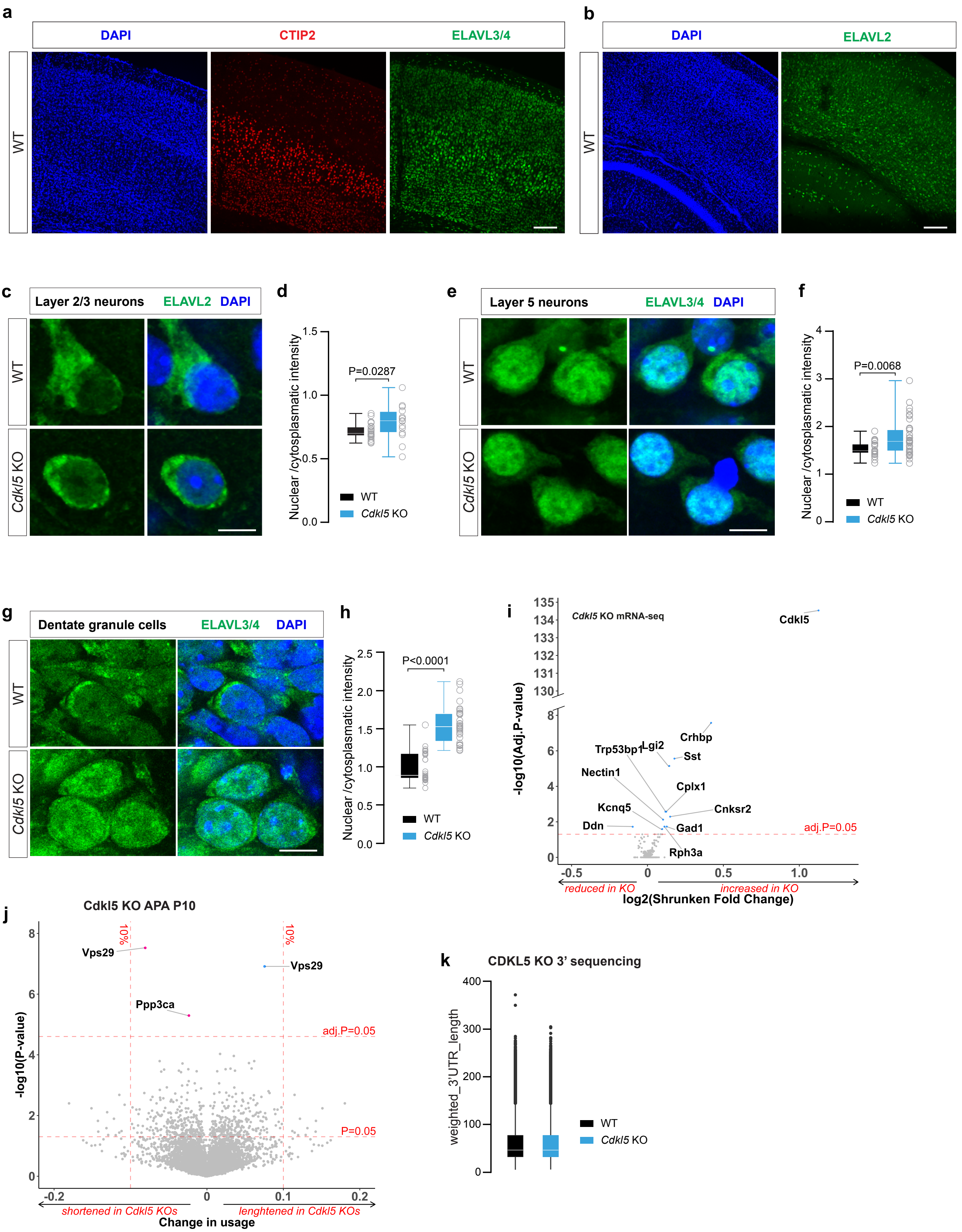

a
DAPI
CTIP2
ELAVL3/4
WT
b
DAPI
ELAVL2
WT
c
Layer 2/3 neurons
ELAVL2
DAPI
WT
Cdkl5 KO
d
Nuclear /cytosplasmatic intensity
P=0.0287
WT
Cdkl5 KO
e
Layer 5 neurons
ELAVL3/4
DAPI
WT
Cdkl5 KO
f
Nuclear /cytosplasmatic intensity
P=0.0068
WT
Cdkl5 KO
g
Dentate granule cells
ELAVL3/4
DAPI
WT
Cdkl5 KO
h
Nuclear /cytosplasmatic intensity
P<0.0001
WT
Cdkl5 KO
i
Cdkl5 KO mRNA-seq
Cdkl5
-log10(Adj.P-value)
Crhbp
Trp53bp1
Lgi2
Sst
Nectin1
Cplx1
Kcnq5
Cnksr2
Ddn
Gad1
adj.P=0.05
Rph3a
reduced in KO
increased in KO
log2(Shrunken Fold Change)
j
Cdkl5 KO APA P10
-log10(P-value)
10%
Vps29
Ppp3ca
adj.P=0.05
P=0.05
shortened in Cdkl5 KOs
lenghtened in Cdkl5 KOs
Change in usage
k
CDKL5 KO 3' sequencing
weighted_3'UTR_length
WT
Cdkl5 KO

**Supplementary Fig. 3: Localization of nELAVL proteins in mouse brain.**

**a, b** Immunostainings for endogenous ELAVL3/4 (**a**) and ELAVL2 (**b**) in WT brains show expected cortical distribution. Scale bar = 200µm. **c** High-magnification view of layer 2/3 pyramidal neurons from WT and *Cdkl5* knockout mice at P10 stained for ELAVL2. The scale bar is 10 µm. **d** Quantification of ELAVL2 immunofluorescence. n=22-36 neurons/group (Two-sided unpaired t-test, WT=1.538, SEM=0.03392; *Cdkl5* KO=1.776, SEM=0.06269). **e** Higher-magnification view of layer 5 pyramidal neurons from WT and *Cdkl5* knockout mice at P10 stained for ELAVL3/4. **f** Graph shows the ratio between nuclear/cytoplasmic immunofluorescence intensity for ELAVL3/4. n=15-24 neurons/group. Scale bar=10 µm (Two-sided unpaired t-test was used to assess statistical significance, WT=0.6170, SEM=0.01240; *Cdkl5* KO=0.6873, SEM=0.03384). **g** Dentate granule cells of hippocampus are analyzed upon immunostaining with ELAVL3/4 antibody. Scale bar=10 µm. **h** Nuclear to cytoplamic ratio is increased in *Cdkl5* KO mice. n=29-32 neurons/group (Two-sided unpaired t-test was used to assess statistical significance, WT=1.000, SEM=0.03711; *Cdkl5* KO=1.558, SEM=0.04308). **i** Bulk mRNA sequencing comparing WT and *Cdkl5* KO mice. *Cdkl5* mRNA is increased in *Cdkl5* KO brains, y axis is discontinous due to the adjusted p-value for *Cdkl5* mRNA being highly significant. Even though *Cdkl5* mRNA is increased, mRNA contains the exon 4 deletion. **j** Volcano plot of P10 alternative poly-adenylation (APA) analysis comparing *Cdkl5* KO vs WT using 3’RNA-seq. Significance criteria are adj.p-value<0.05 and change in usage>10%. n=5 mouse brains per group. **k** Weighted 3’UTR length analysis using 3’RNA-sequencing approach shows no differences between WT and *Cdkl5* KO animals.
