## Supplementary Figure 4 for "CDKL5 phosphorylates neuronal ELAVL proteins to promote mRNA binding, protein synthesis and visual cortex development"

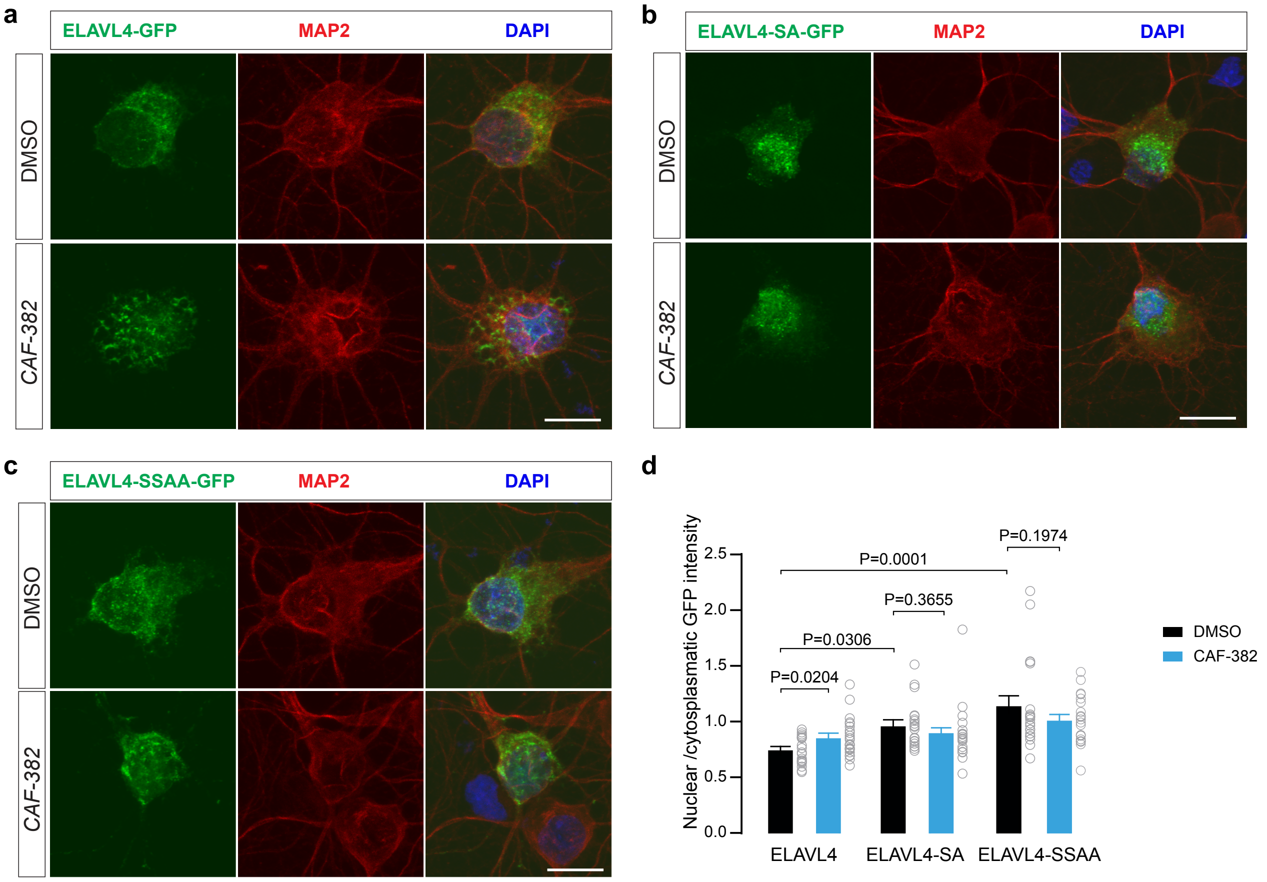


**Supplementary Fig. 4: nELAVL localization in nucleus and cytoplasm upon treatment with CDKL5 inhibitor.**

**a-c** Distribution of GFP-tagged ELAVL4 WT (**a**), ELAVL4-SA (S131A) (**b**) and ELAVL4-SSAA (S130A and S131A) (**c**) in neurons treated with DMSO or CAF-382 (CDKL5 inhibitor) for 4 hours. **d** Quantification from (**a**), (**b**) and (**c**). DMSO, n=19-21 neurons; CAF-382, n=19-23 neurons; SEM error bars, (ANOVA followed by Dunnett's multiple comparisons test, ELAVL4-GFP (Control)=0.7719, SEM=0.02968; ELAVL4-GFP (CAF-382)=0.8900, SEM=0.03711; ELAVL4-SA-GFP (Control)=1.006, SEM=0.04842; ELAVL4-SA-GFP (CAF-382)=0.9340, SEM=0.06242; ELAVL4-SSAA-GFP (Control)=1.183, SEM=0.09033; ELAVL4-SSAA-GFP (CAF-382)=1.049, SEM=0.04918). Scale bars: 20 µm.
