## Supplementary Figure 5 for "CDKL5 phosphorylates neuronal ELAVL proteins to promote mRNA binding, protein synthesis and visual cortex development"

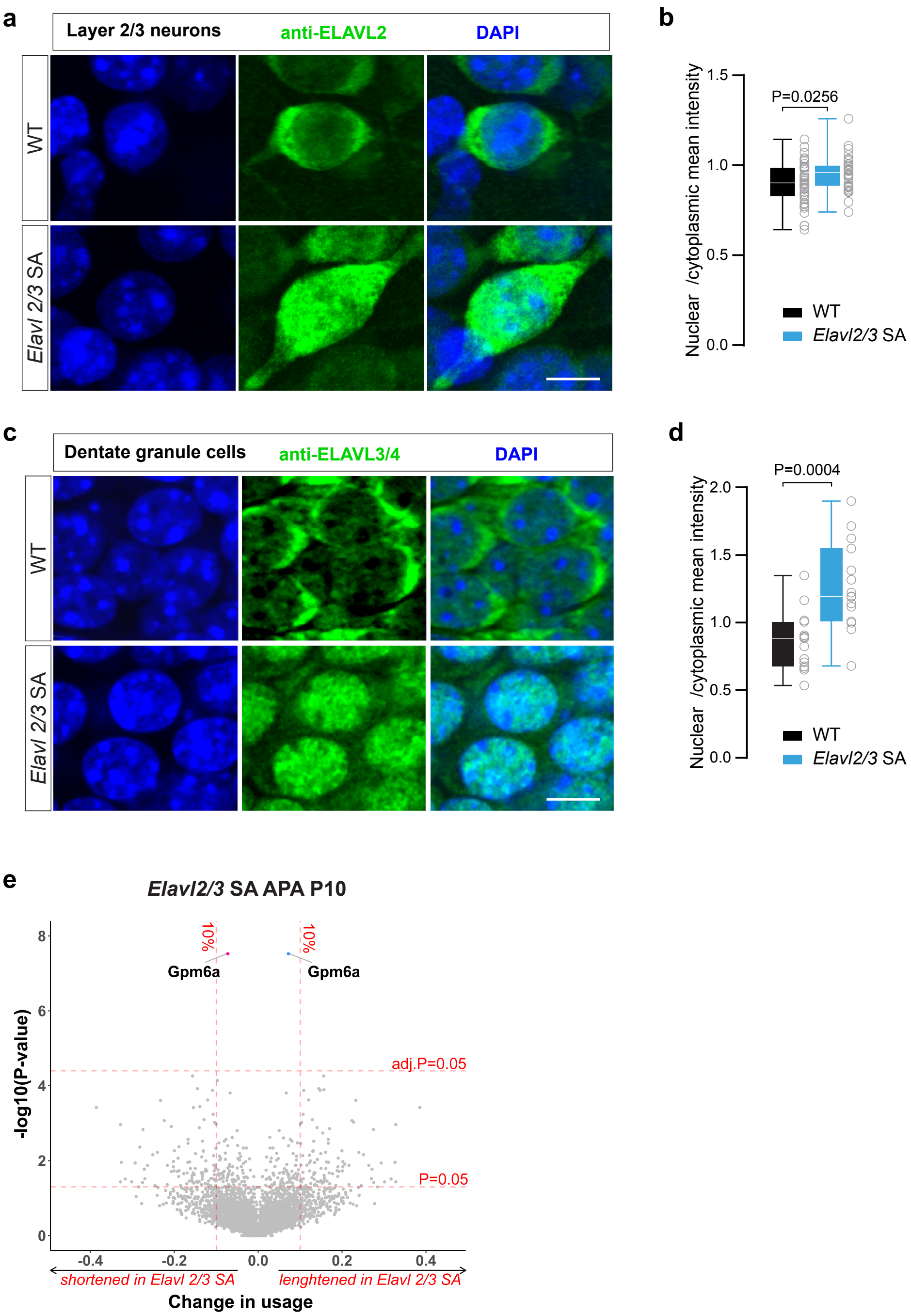


**Supplementary Fig. 5: Proteomic and transcriptomic analysis of *Elavl2/3* SA phosphomutant mice.**

**a** High-magnification view of layer 2/3 pyramidal neurons from WT and *Elavl2/3* phosphomutant mice at P15 immunostained with ELAVL2 antibody. Scale bar=10 µm. **b** Quantification of the ratio between nuclear and cytoplasmic immunofluorescence signals of ELAVL2 in P15 *Elavl2/3* phosphomutant and WT mice. n=38-39 neurons per group. Two-sided Student's t test was used to assess statistical significance (WT=0.8981, SEM=0.01858; *Elavl 2/3* SA=0.9524, SEM=0.01486). **c** Endogenous ELAVL3/4 immunostainings in dentate granule neurons from WT and *Elavl2/3* phosphomutant mice. **d** Graph shows the ratio between nuclear and cytoplasmic immunofluorescence signal for ELAV3/4. n=15 neurons/group. Scale bar=10 µm. Two-sided Student's t test was used to assess statistical significance (WT=0.8601, SEM=0.05550; *Elavl 2/3* SA=1.267, SEM=0.08343). **e** Volcano plot of P10 alternative poly-adenylation (APA) analysis comparing *Elavl2/3* SA vs WT using 3'RNA-seq. Significance criteria are adj.p-value<0.05 and change in usage>10%. n=5 mouse brains per group.
