## Supplementary Figure 6 for "CDKL5 phosphorylates neuronal ELAVL proteins to promote mRNA binding, protein synthesis and visual cortex development"

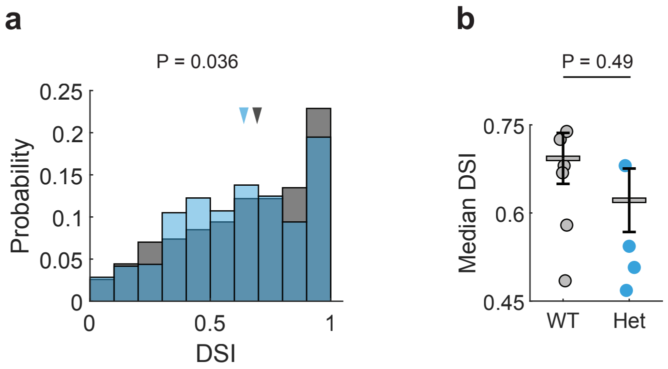


**Supplementary Fig. 6: DSI analysis.**

**a** Distribution of direction selectivity index (DSI) for visual neurons of WT and Het animals. Triangles indicate the median of the distribution. **b** Median DSI for individual recordings. P-values were calculated using Mann–Whitney U test.
